# Autism-risk gene mutations convergently disrupt sexually dimorphic oxytocin circuits to lower social engagement

**DOI:** 10.64898/2026.08.27.747579

**Authors:** A. Patwardhan, A. Moslemian, E. Tse, V. Grinevich, K.Y Choe

**Author notes:** Corresponding author: Katrina Choe.

## Abstract

Autism arises from diverse genetic risk factors, yet how they converge to produce core symptoms and contribute to its sex bias remains unestablished. Oxytocin increases sociability in multiple murine autism models, presenting an opportunity to identify a potentially shared mechanistic basis across etiologies. Here we show that spontaneous social investigation triggers overlapping patterns of aberrant functional connectivity across social and sensory brain regions in two knockout (KO) mouse models, which are rescued by oxytocin. We also report that, during social investigation, wildtype mice exhibit sexually dimorphic oxytocin release and neuronal activity dynamics in the nucleus accumbens and the amygdala. These patterns are disrupted in both KO models, but can be restored by sex- and circuit-specific stimulation of endogenous oxytocin release, accompanied by enhanced social engagement. These findings identify impaired oxytocin recruitment of sexually dimorphic social circuits as a convergent consequence of autism-risk gene mutations that may underlie low sociability.

## Introduction

Autism spectrum disorder (ASD) is diagnosed 3-4 times more frequently in boys than in girls^1^. While biological factors are estimated to be major contributors to this difference in prevalence^2–4^, another contributing factor is the “milder” presentation of behavioral symptoms in females that often leads to underdiagnosis, especially in the social domain^5–7^. While males and females have been shown to utilize largely overlapping brain networks to support social behavior, sex differences in functional recruitment of these circuits have been documented^8–10^ with proposed contributions to sex-specific social behavioral expression. Despite the major sex differences in ASD prevalence and behavioral presentation, the extent to which sexually dimorphic neural mechanisms contribute to behavioral symptom differences in males and females remains largely unknown.

ASD has a complex genetic etiology, featuring over 1,000 candidate genes discovered through large-scale genome-wide sequencing efforts over the last two decades^11^. Despite this genetic heterogeneity, accumulating experimental evidence suggests that diverse genetic disturbances converge on common neurobiological pathways that contribute to shared behavioral features of ASD^12–14^. In parallel with the rapidly growing list of ASD-risk genes, the last two decades have witnessed an expanding collection of mouse models that feature high-risk genetic disruptions for ASD, many of which exhibit reproducible ASD-like behavioral disruptions^11,15–18^. Rigorous investigation of these models has revealed mechanistic links between each genetic perturbation and their behavioral phenotypes; however, due to the large number of models and often variable behavioral phenotypes, establishing a systems-level consensus on the neural mechanisms that underlie ASD-like behaviors still represents a formidable challenge. Importantly, there is a bias toward males in studies with mouse models of ASD^4^, preventing the discovery of potential sex-specific neural pathways with behavioral implications.

Two of the most well-characterized genetic models of ASD are *Cntnap2* knockout (KO)^19–23^ and *Fmr1* KO^24,25^ mice, both modelling loss-of-function mutations associated with syndromic forms of ASD. Remarkably, both models feature evidence of central oxytocin (OXT) deficiency, as well as a rescue of social phenotypes after OXT administration^26–28^. Together with the well-established neuromodulatory influences of OXT on various forms of social behavior^29–34^, these observations raise the possibility that these distinct gene mutations convergently disrupt the function of the central OXT system to lower the sociability in both KO models. A major unanswered question is whether the central OXT deficiency in the two models disrupts social behavior through shared or distinct circuit mechanisms, and whether they differ by sex. Defining these pathways is crucial for understanding the interplay between ASD-risk gene mutations and potentially shared social circuit mechanisms, which may be sexually dimorphic.

Here we show, in both *Cntnap2* KO and *Fmr1* KO mice, a shared phenotype of reduced social engagement duration with subtle sex differences. This behavioral phenotype is associated with convergent disruptions in activity and functional connectivity within the social salience network (SSN), an interconnected collection of brain regions known to play a crucial role in evaluating socially relevant sensory signals and modulating social engagement^35,36^. Chemogenetically stimulating endogenous OXT release reverses these phenotypes in both models. We further demonstrate, in WT mice, that social investigation triggers sexually dimorphic increases in neuronal activity as well as OXT levels in two key social regions: the nucleus accumbens shell (NAcSh) in males, and the lateral subdivision of the central amygdala (CeL) in females. In both KO models, social investigation instead reduced NAcSh neuronal activity in males and CeL neuronal activity in females, coupled with a lack of OXT release within respective regions. Finally, sex-specific rescue of endogenous OXT release in appropriate circuit pathways corrected both neuronal activity and social behavioral phenotypes in males and females. Taken together, these data reveal that OXT modulates social circuits in a sex-dependent manner to dictate the length of social engagement. Furthermore, our results provide strong evidence for OXT modulation of sexually dimorphic social circuits as a convergent mechanism underlying social behavior disruptions caused by distinct ASD-risk gene mutations.

## Results

### Sex- and genotype-specific differences in social investigation

We first sought to systematically compare the social investigation phenotypes in both male and female *Cntnap2*^-/-^ and *Fmr1^-/-,-/Y^* mice using an ethologically relevant paradigm. Conflicting reports of social phenotypes in KO mouse models of ASD-risk gene mutations are quite common^37–41^, including sex differences in *Fmr1^-/-,-/Y^*^42,43^ and *Cntnap2*^-/-^ mice^38,44^. A large proportion of studies utilize the 3-chambered social approach test, which was designed to specifically test a mouse’s preference for a wire cage containing a stimulus mouse vs. another wire cage that is either empty or contains a novel object^45^. However, the lack of direct social interaction in this test prevents the expression of nuanced social behavior^46,47^, and is of limited utility for comparisons of interaction duration^16^, a parameter previously shown to be modulated by OXTergic signalling^48^. Therefore, to more comprehensively examine OXT-relevant social behavior patterns across sexes and genotypes, we examined free social investigation between a pair of unfamiliar mice in a home cage setting, a well-established and ethologically relevant paradigm that has reliably demonstrated sociability differences in ASD models^19,28,38^.

Adult experimental mice (*Cntnap2*^-/-^, *Fmr1*^-/-,-/Y^, or their WT littermates) were paired with juvenile (3-5 weeks old) WT mice and allowed to freely interact in a neutral home cage setting for 10 minutes, similar to commonly used social interaction paradigms designed to promote non-aggressive social investigation between unfamiliar partners in both males and females (**Fig.1a**)^19,28^. Consistent with previous reports^19,28^, both male and female *Cntnap2^-/-^*mice exhibited a reduced total duration of social investigation relative to sex-matched *Cntnap2*^+/+^ (WT) littermate controls (q_females_=0.03, q_males_=0.0012; Two-way ANOVA; **Table.1, Fig.1b**). Notably, within the *Cntnap2*^-/-^ group, females showed a significantly lower total duration of investigation than males, indicating a sex difference (q=0.042; **Fig.1b**). However, in comparison with sex-matched *Cntnap2*^+/+^ mice, *Cntnap2*^-/-^ males showed a larger reduction in the total duration of social investigation than *Cntnap2*^-/-^ females (q=0.0008; **Fig.1b**). This apparent disparity in findings is due to *Cntnap2*^+/+^ males showing higher total social investigation duration than *Cntnap2*^+/+^ females (q=0.0016; **Fig.1b**), consistent with previously reported results^49^. In contrast, *Fmr1*^−/Y^ (male) mice exhibited a reduced total duration of investigation (q<0.0001; two-way ANOVA; **Table.1**; **Fig.1b**), but not *Fmr1*^+/+^ (female) mice (q=0.149; **Fig.1b**). Consistent with the above observation in the *Cntnap2* cohort, a sex difference was observed between *Fmr1*^+/Y^ (male) and *Fmr1*^+/+^ (female) mice, with females displaying shorter total duration of investigation than males (p=0.009; **Fig.1b**). Together, these results demonstrate sex differences in the total duration of social investigation in WT mice, as well as reduced social investigation in both male and female *Cntnap2* KO mice but only in male *Fmr1* KO mice. Despite these model-specific effects, both KO models retain sex differences in the duration of social investigation, consistent with clinical observations that distinct genetic risk factors can give rise to sex-dependent differences in behavioral presentation^5–7^.

Next, we examined in each mouse line whether these differences were driven by the number of social bouts (indicating reduced initiation of social investigation) or the duration of each social bout (indicating reduced maintenance of social investigation). In *Cntnap2*^-/-^ mice, we found no differences across conditions in the total number of social bouts (Two**-**way ANOVA; **Fig.1c, Table.1**). Rather, both male and female *Cntnap2*^-/-^ mice had a shorter mean duration of individual social bouts than their WT littermates (Two-way ANOVA, **Fig**.**1d, Table.1**). Similarly, *Fmr1*^-/Y^ (male) mice showed no difference in the number of social bouts (**Fig**.**1c**) but shorter duration of individual social bouts (**Fig.1d**). Surprisingly, *Fmr1*^-/-^ (female) mice, which showed shorter individual social bouts compared to *Fmr1^+/+^*mice (q<0.0001; **Fig.1d**), also showed a greater number of social bouts compared to their WT littermates (q=0.019; **Fig.1c**), explaining the absence of a social phenotype in our earlier comparison of total social investigation duration (**Fig.1b**). In summary, we demonstrate in both *Cntnap2^-/-^*and *Fmr1^-/-,-/Y^*mice, shorter durations of individual social bouts as the primary shared social phenotype in both sexes, with sex differences in the social bout duration within each genotype. This premature termination of social investigation in KO mice may be symptomatic of insufficient social reward signaling^50^.

### Endogenous evoked release of OXT convergently rescues social behavior phenotypes in *Cntnap2* and *Fmr1* KO mice

We and others have previously demonstrated that experimental stimulation of endogenous OXT release reliably rescues the social behavior phenotypes in *Cntnap2^-/-^* mice^19,28^; however, it has not been examined whether this behavioral rescue specifically increases the duration or number of social investigation bouts in *Cntnap2^-/-^* mice or any other KO models of ASD-risk gene mutations. To directly test this, we chemogenetically activated paraventricular (PVN)-OXT neurons using the excitatory Designer Receptors Exclusively Activated by Designer Drugs (DREADD) hM3Dq, to stimulate endogenous central OXT release^19,28,51^ prior to subjecting mice to the home cage social investigation (**Fig.1e**). We found that clozapine-n-oxide (CNO) activation of hM3Dq in PVN-OXT neurons significantly increased social investigation time in both *Cntnap2^-/-^*and *Fmr1^-/-,-/Y^* mice compared to CNO administered control mice (Oxt-Venus; p*_Cntnap2_*=0.033, p*_Fmr1_*=0.037; unpaired t-test), an effect that was driven by the mean duration (p*_Cntnap2_*=0.0091, p*_Fmr1_*=0.0093; unpaired t-test), but not the number (p*_Cntnap2_*=0.31, p*_Fmr1_*=0.49, unpaired t-test) of investigation bouts (**Fig.1f,g**). Additionally, we found that hM3Dq activation decreased the interval between consecutive social bouts in *Cntnap2^-/-^* and *Fmr1^-/Y^* (male) mice (p*_Cntnap2_*=0.037, p*_Fmr1_*=0.012; Two sample Kolmogorov-Smirnov test; **Fig.1h,i**), indicating that subsequent investigation bouts occur more quickly. This effect was absent in *Fmr1^-/-^* females (**Fig.1i**). In contrast, hM3Dq activation lengthened the duration of social bouts in males and female KO mice (**Fig.1h,i**). Collectively, these data suggest that chemogenetic stimulation of endogenous central OXT release ameliorates the social behavior phenotype by lengthening the duration of social investigation bouts in both KO mouse lines.

### PVN-OXT neuron activation alters social investigation-associated regional activity and functional connectivity within SSN of KO mice

Our previous work demonstrated that exogenous OXT administration, in the absence of a social context, activates a network of brain regions with established roles in social behavior in *Cntnap2^-/-^* mice^19^. We further showed that chemogenetically stimulating endogenous central OXT release, also in the absence of a social context, robustly increased neuronal activity in the nucleus accumbens (NAc). This is directly in line with the proposed role of the NAc as a hub of the SSN, a collection of interconnected brain regions thought to encode the salience and rewarding value of social stimuli and to be strongly modulated by OXT^35,52^. However, it is likely that activity changes in other brain regions that are also crucial for social investigation went undetected due to the absence of social context in our previous study. Therefore, we sought to comprehensively map SSN activity patterns associated with social investigation in *Cntnap2^-/-^* mice, determine how these patterns are altered by stimulated central OXT release, and test whether similar alterations are present in *Fmr1^-/-,-/Y^*mice.

To perform a systematic assessment of regional activity changes within the SSN with minimal data loss, we used a whole-brain c-Fos activity mapping pipeline combining SHIELD tissue clearing/c-Fos+ immunolabelling and lightsheet imaging (**Fig.2a**)^53,54^. We collected the brains of *Cntnap2^-/-^* and *Fmr1^-/-,-/Y^*mice after they socially investigated novel mice in the home cage setting, then processed them for whole-brain SHIELD-tissue clearing/c-Fos+ immunolabelling. Lightsheet images from these brains were then analyzed to quantify c-Fos+ cell counts from 14 SSN brain regions, which were compared between groups (**Fig.2a,b**). First, we examined whether specific brain areas within the SSN showed significant increases or decreases in c-Fos+ cell counts due to chemogenetic stimulation of endogenous OXT release during social investigation. Compared to Oxt-Venus controls, Oxt-hM3Dq groups in *Cntnap2^-/-^* and *Fmr1^-/-,-/Y^*cohorts showed a significantly decreased number of c-Fos+ cells in the hippocampal formation (HPF) and somatosensory cortex (SS) (q*_Cntnap2_*-_HPF_<0.001, q*_Fmr1_*-_HPF_<0.001, q*_Cntnap2_*_-SS_=0.0017, q*_Fmr1_*_-SS_=0.012; one-way ANOVA with FDR correction; **Fig.2c, Table.2a**). Additionally, we observed non-significant increases in the number of c-Fos+ cells within the infralimbic area (ILA), PVN, NAc, CeA, and the anterior olfactory nucleus (AON), all of which showed striking consistency between the two KO models.

Next, we examined whether chemogenetic stimulation modified functional connectivity across the SSN by computing Pearson’s correlation coefficient (*r*) between c-Fos+ cell counts across pairs of SSN regions, and comparing the pairwise *r* values between control and hM3Dq-treated KO mice. In the control group, we found that the NAc either showed a lack of correlation or negative correlation with a large number of other SSN regions in both KO mdodels, including HPF, BLA, VIS, and AUD in *Cntnap2^-/-^* mice and HPF, MeA, BLA, VIS, AUD, and SS in *Fmr1^-/-,-/Y^* mice (**Fig.2e,f**). Strikingly, hM3Dq activation of PVN*-*OXT neurons either eliminated these negative correlations (HPF, BLA, and AUD in *Cntnap2^-/-^* mice, and HPF, BLA, AUD, SS in *Fmr1^-/-,-/Y^* mice), or flipped them to positive correlations (VIS in *Fmr1^-/-,-/Y^* mice) in both KO models (**Fig.2e,f, Table.2b**). The most notable connectivity change, however, was observed in the sensory processing areas and the subnuclei of the amygdala (BLA, MeA, CeA), where mice in the control condition showed strong positive correlations with sensory areas (SS, AON, AUD, VIS) that became decoupled in the hM3Dq condition (**Fig.2e,f, Table.2b**). This effect was consistent across the two genotypes, suggesting these circuits as potential sites of model convergence. There were also a few notable differences in connectivity observed between the KO models. In the control condition, the AON was positively correlated with the amygdala in *Cntnap2^-/-^*mice (r_CeA_=0.69, r_MeA_=0.82, r_BLA_=0.95; Pearson Correlation; **Fig.2e,f, Table.2b**), whereas in *Fmr1^-/-,-/Y^*mice the AON was anticorrelated with the amygdala (r_CeA_=-0.17, r_MeA_=-0.62, r_BLA_=-0.49). In the hM3Dq treatment condition, however, the AON showed similar functional connectivity patterns in both models. Furthermore, in the hM3Dq treatment condition, the medial amygdala (MeA) was positively correlated with the ILA and PVN in *Cntnap2^-/-^* mice (r_ILA_=0.47, r_PVN_=0.67), whereas in *Fmr1^-/-,-/Y^* mice, it was anticorrelated (r_ILA_=-0.20, r_PVN_=-0.19). Together, these results demonstrate that hM3Dq-mediated stimulation of endogenous OXT release promotes social investigation-associated coordination of activity in the ILA, PVN, NAc and CeA in KO mice in a model convergent manner.

While our above data demonstrate that chemogenetic activation of PVN-OXT neurons can activate specific SSN nodes in both mouse lines in a consistent manner (**Fig.2**), it is still unclear whether and which specific nodes of the SSN are most relevant to the social investigation phenotypes and the hM3Dq-induced behavioral rescue we observed in the two KO models (**Fig.1**). Therefore, we correlated across mice the mean duration or number of social bouts in each mouse with the number of c-Fos+ cells from individual SSN regions. We found that regional activity levels of the ILA, PVN, NAc, CeA and SS are strongly correlated with mean duration of individual social bouts in both *Cntnap2^-/-^* and *Fmr1^-/-,-/Y^* mice (**Fig.2g,h**; **Table.2b)**. The number of c-Fos+ cells in all of the above regions was positively correlated with the mean duration of social investigation bouts, except for SS, which showed a negative correlation (**Fig.2g,h**). Interestingly, across genotypes, data points representing animals in the hM3Dq-treated cohort clustered at the top-right or bottom-right depending on the regions that showed positive or negative correlations with the mean duration of investigation bouts (**SFig.1**). In contrast, the number of social investigation bouts was not correlated with activity in the above-mentioned regions, but was instead correlated only with the anterior cingulate cortex (ACC) (**Fig.2, SFig.2**). These findings suggest that enhanced OXT signaling differentially modulates the activity of individual SSN regions, producing region-specific activity patterns associated with prolonged social investigation bouts. We further identify with these data the ILA, PVN, NAc, and CeA as OXT-sensitive key SSN regions whose activity levels are positively associated with sustained social engagement.

**Figure 1.**
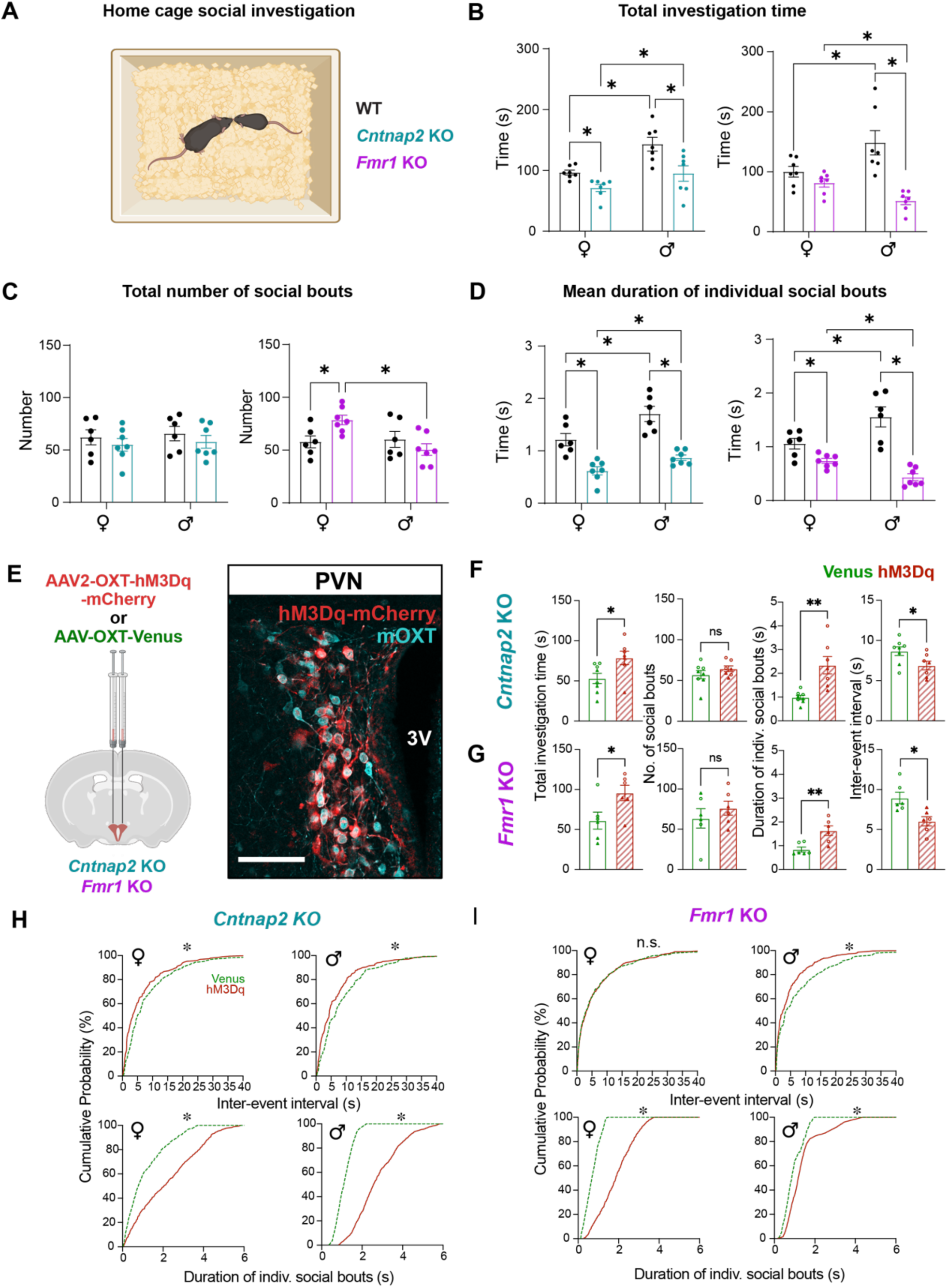
Shorter duration of individual social bouts is a shared behavioral feature of *Cntnap2* and *Fmr1* KO mice, rescued by stimulating endogenous OXT release. a. Behavior schematic. **b-d.** Sex stratified comparisons between *Cntnap2* (green) and *Fmr1* KO (purple) mice with their WT littermates (black). Total duration (**b**), total number (**c**) and (**d**) mean duration of social investigation bouts. Two-way ANOVA with FDR correction for multiple comparisons. **e.** Validation of hM3Dq (red) expression in PVN colocalized (white) with immunolabeled OXT neurons (cyan). *Left*, Schematic. *Right,* OXT-hM3Dq or OXT-Venus (control) injection in PVN. 3V, 3^rd^ ventricle. Scale bar=100μm. **f-g.** Comparisons of total investigation time and mean duration of investigation bouts between hM3Dq (red) and Venus control (green) conditions in *Cntnap2* and *Fmr1* KO mice. Two-sample t-test. **h-i**. Sex-stratified cumulative probability plots comparing time intervals between consecutive social bouts (top) and mean duration of social bouts (bottom) between hM3Dq (red) and Venus control (green) conditions in *Cntnap2* (**h**) and *Fmr1* (**i**) KO mice. Kolmogorov-Smirnov test. *P<0.05, **P<0.01, ns=not significant.

**Figure 2.**
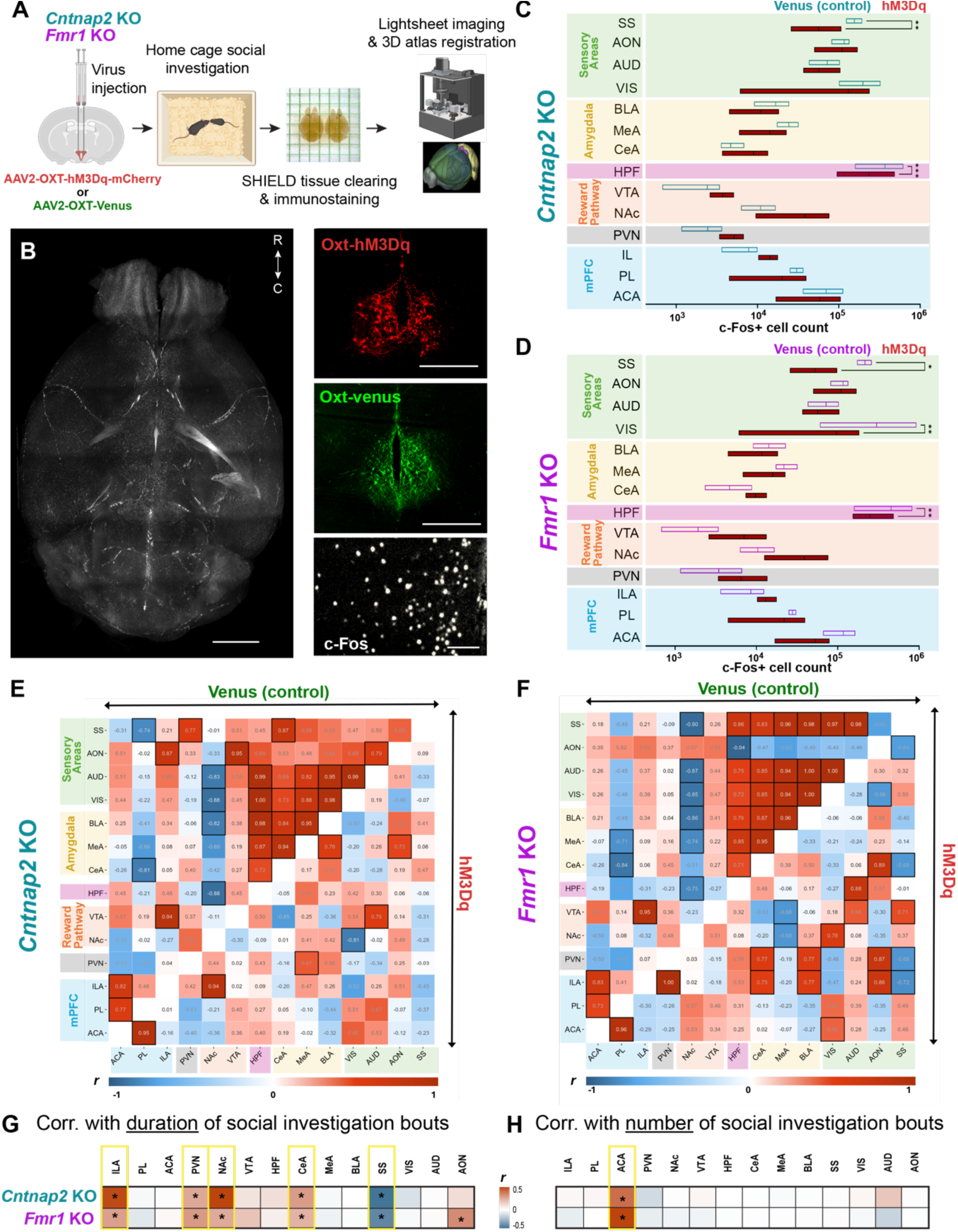
DREADD-mediated PVN-OXT neuronal activation alters social investigation-associated regional activity and functional connectivity within SSN of KO mice. **a.** Schematic: experimental strategy (atlas parcellation image^116^). b. *Left*, representative lightsheet image of whole brain c-Fos labeling. Scale bar=2mm. *Right,* OXT neurons labeled with hM3Dq (red, top). Scale bar=500μm, Venus (green, middle). Scale bar=500μm. Zoomed-in image showing cortical c-Fos+ cells (bottom). Scale bar=40μm. **c-d.** Comparisons of c-Fos+ cell counts in SSN regions across treatment conditions and genotypes (c. *Cntnap2* KO, d. *Fmr1* KO). One-way ANOVA with FDR correction. *P<0.05, **P<0.01, ***P<0.001. **e-f.** Cross-correlation heatmaps comparing network-level activity in the Control (*top left*) vs hM3Dq (*bottom right*) groups across genotypes (e. *Cntnap2* KO, f. *Fmr1* KO). Black squares indicate non-zero Pearson correlations with P<0.05. **g-h.** Heatmaps showing Pearson correlations between c-Fos+ cell counts and mean duration (**g**) and total number (**h**) of social investigation bouts in SSN regions. Yellow boxes represent Pearson correlations with P<0.05 that are common in *Cntnap2* (top) and *Fmr1* (bottom) KO mice.

### Cntnap2 or Fmr1 KO has sex-specific effects on social investigation-induced changes in c-Fos-positive cell density in major SSN nodes

The chemogenetically activated SSN activity patterns we reported above reveal the brain areas that potentially mediate the OXT-induced social rescue in both KO models, but it is unknown whether similar patterns are naturally elicited by social investigation in WT mice with intact central OXT system function. To directly test this, we examined coordinated activity among ILA, PVN, NAc, and CeA, identified above as key SSN regions for OXT-induced social rescue, in the presence or absence of social stimuli in both KO models and their WT littermates. First, we examined the coordinated patterns of activity across these 4 regions in *Cntnap2^-/-^* and *Fmr1^-/-,-/Y^*mice and WT controls, split into two groups: one group was placed in a home cage together with a juvenile sex-matched WT stimulus mouse (identical to the previous home cage social investigation experiments), and the other group was placed alone in a home cage to serve as controls (**Fig.3a**). Post-behavior, we quantified c-Fos+ cell density in each of the 4 brain regions from immunostained brain sections imaged with a slidescanner. In the home cage condition without a social stimulus, all mice showed largely uncoordinated activity across the 4 regions (**Fig.3b**). In the home cage social investigation group, WT mice showed strong positive correlations in activity between all pairs of examined regions, suggesting increased functional connectivity across the 4 SSN nodes (**Fig.3b, Table.3**). Strikingly, both *Cntnap2^-/-^* and *Fmr1^-/-,-/Y^* mice showed no significant correlations in activity within this subnetwork post-social investigation (**Fig.3b, Table.3**).

To examine whether the changes in activity in the 4 SSN nodes was accompanied by a strengthening of functional connectivity across these regions in WT and to compare with KO mice, we quantified social investigation-triggered changes in c-Fos+ cell density within individual SSN nodes of *Cntnap2^-/-^* and *Fmr1^-/-,-/Y^* mice and WT littermates. Because previous studies have shown sex differences in neural pathways that encode and process social information in WT mice^55–58^, we conducted sex-stratified comparisons of c-Fos+ cell density. Consistent with previous reports^59–62^, we found that social investigation caused an increase in c-Fos+ cell density in the ILA and PVN of WT mice (**Fig.3c,d, Table.3**). However, this canonical increase in activity was not observed in the ILA in *Cntnap2^-/-^*and *Fmr1^-/Y,-/-^* mice. More interestingly, this analysis revealed several sex- and region-specific activity phenotypes that were consistent across *Cntnap2^-/-^* and *Fmr1^-/-,-/Y^* mice. Both *Cntnap2*^-/-^ and *Fmr1^-/Y^* males showed decreased PVN c-Fos+ cell density after social investigation (q*_Cntnap2_*_,male_=0.0117, q*_Fmr1_*_,male_<0.001; **Fig.3c,d**), while female *Cntnap2*^-/-^ and *Fmr1^-/-^* mice did not show a significant change after social investigation (q*_Cntnap2_*_,female_=0.104, q*_Fmr1_*_,female_=0.062; **Fig.3c,d**).

This sex difference in social investigation-induced PVN activity could have sex-specific implications for the activity of downstream regions such as the NAc and CeA. In fact, we observed a stark sex difference in social investigation-triggered activity in both regions of WT mice, as well as sex-specific activity phenotypes in KO mice. In the NAc, we observed increased c-Fos+ cell density in WT males after social investigation (q*_Cntnap2_*_,male_=0.0013, q*_Fmr1_*_,male_<0.001; **Fig.3c,d**), whereas both *Cntnap2*^-/-^ and *Fmr1^-/Y^* males showed decreased NAc c-Fos+ cell density (q*_Cntnap2_*_,male_=0.002, q*_Fmr1_*_,male_=0.002; **Fig.3c,d**). In contrast, social investigation had no effect on NAc activity in WT, *Cntnap2*^-/-^, and *Fmr1^-/-^* females (q*_Cntnap2_*_,female_=0.4402, q*_Fmr1_*_,female_=0.479; **Fig.3c,d**). These surprising results could indicate a male-biased role of the NAc in modulating free social investigation. In the CeA, WT females, but not males, showed increased c-Fos+ cell density after social investigation (q*_Cntnap2+/+,_*_female_=0.002, q*_Fmr1+/+,_*_female_=0.004; **Fig.3c,d**). In contrast, social investigation reduced CeA c-Fos+ cell density in both *Cntnap2*^-/-^ and *Fmr1^-/-^* females (q*_Cntnap2_*_,female_=0.004, q*_Fmr1_*_,female_=0.003; **Fig. 3c,d**). WT, *Cntnap2*^-/-^, and *Fmr1^-/Y^* males showed no change in the c-Fos+ cell density after social investigation (q*_Cntnap2+/+,_*_male_=0.158, q*_Cntnap2-/-_* _,male_=0.214, q*_Fmr1+/Y,_*_male_=0.134, q*_Fmr1-/Y_*_,male_=0.332; **Fig.3c,d**), suggesting a female-specific role of the CeA WT social investigation.

**Figure 3.**
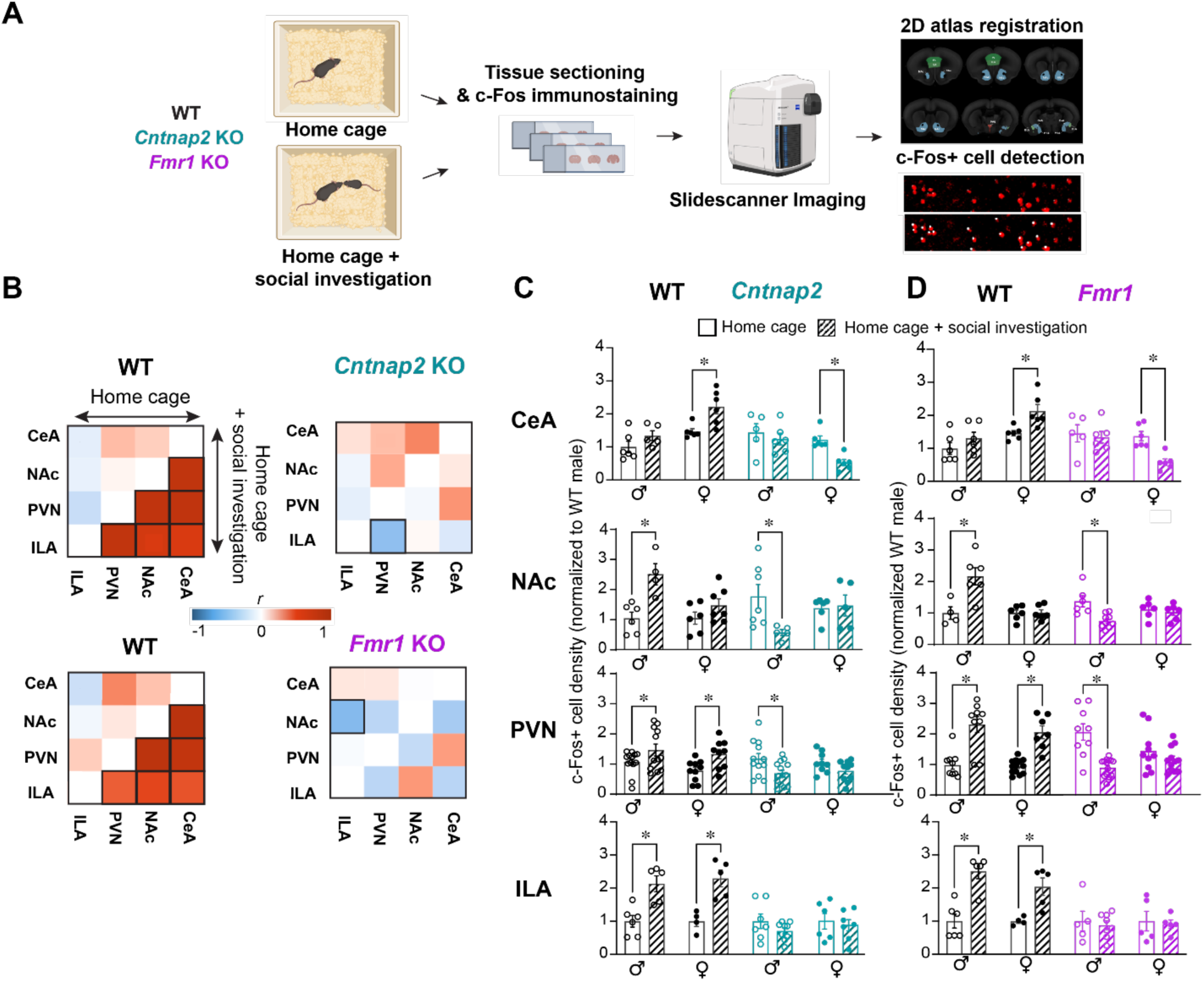
*Cntnap2* or *Fmr1* KO has sex-specific effects on social investigation-induced changes in c-Fos-positive cell density in 4 key SSN nodes. a. Schematic: experimental design. **b.** Cross-correlation heatmaps comparing activity in CeA, NAc, PVN and ILA after home cage (top left of heatmap square) vs home cage+social (bottom right of heatmap square) across genotypes: *Cntnap2* (top right) and *Fmr1* KO (bottom right) mice and their WT littermates (left). Squares with black borders represent non-zero Pearson correlations with P<0.05. **c-d.** WT baseline-normalized c-Fos+ cell density changes in the CeA, NAc, PVN and ILA in *Cntnap2* (green) and *Fmr1* KO (purple) mice and their WT littermates (black). *P<0.05.

It is important to note that, given inhibitory circuit architecture within the CeA, the lateral (CeL) and medial (CeM) subregions of the CeA largely show inversely correlated activity due to the GABAergic projections from the CeL to the CeM^62^. Indeed, we observed opposite social investigation-induced activity patterns in these subregions (**SFig.3**). Notably, it is the CeL that showed increased activity during social investigation in both WT males and females (q*_Cntnap2+/+,_*_male_<0.0001, q*_Cntnap2+/+,fe_*_male_=0.0002, q*_Fmr1+/Y,_*_male_<0.0001, q*_Fmr1+/+_*_,female_=0.0005; **SFig.3**), consistent with previous findings that the CeL receives the large majority of OXTergic input from the PVN^63,64^. Thus, our data suggest that specific subregions and cell populations within the CeA encode the interplay between OXTergic input and modulation of regional activity during social investigation.

### Cntnap2**^-/-^** and Fmr1**^-/Y^**males exhibit reduced D1-MSN activity and lack social investigation-induced OXT release in the NAcSh

Our above results are highly consistent with our previous demonstration that OXTR activation in the NAcSh is critical for increasing social investigation duration of *Cntnap2^-/-^* mice^19^. Whether social investigation-associated activity in NAcSh neurons is altered in *Cntnap2^-/-^* and *Fmr1^-/-,-/Y^* mice remains unexplored. Within the NAcSh, dopamine 1 receptor (D1R)-expressing medium spiny neurons (D1-MSNs) have been shown to dynamically encode appetitive stimuli including social stimuli^65,66^ and make up a majority of OXTR-expressing cells^58,67^. Therefore, we hypothesized that D1-MSNs in the NAcSh of KO mice exhibit aberrant activity in response to social stimuli and potentially mediate OXT-induced social rescue. To test this, we measured population-level GCaMP8f calcium dynamics in NAcSh D1-MSNs via fiber photometry in KO mice and their WT littermates during social investigation (**Fig.4a**).

Similar to previously reported results, we observed a gradual increase in NAcSh D1-MSN GCaMP8f signal close to the onset of social investigation in male WT mice. In striking contrast, both *Cntnap2*^-/-^ and *Fmr^,-/Y^* males, which had shorter social investigation bouts than WT littermates (unpaired t-tests, p*_Cntnap2, male_*=0.0004, p*_Cntnap2, female_*=0.0233, p*_Fmr1, male_*<0.0001, p*_Fmr1, female_*<0.0001; **SFig.4**), exhibited GCaMP8f signal decays close to the onset of social investigation (**Fig.4b-c,d,f**). Comparisons of social-investigation associated peak values and areas under the curve (AUC) between both KO mice and their WT littermate controls revealed consistent activity changes in opposite directions (p*_Cntnap2_*<0.0001, p*_Fmr1_*<0.0001 for both peak value and AUC comparisons, unpaired t-tests; **Fig.4e,g, Table.4**). Interestingly, in females, NAcSh D1-MSNs did not show any activity alterations during social investigation in WT or KO mice. In all female groups, no peaks were detected in the GCaMP8f signal from NAcSh D1-MSN. Additionally, we report that these sex-specific effects of social investigation on NAcSh D1-MSN activity are strongest in the first investigation bouts at the beginning of the homecage social investigation assay with a steady decay in signal change thereafter (**SFig.4**). Thus, these data reveal a surprising sex-specificity in social circuit engagement (**Fig.4a-i**) that is consistent with our above data from c-Fos analyses (**Fig.3c,d**).

**Figure 4.**
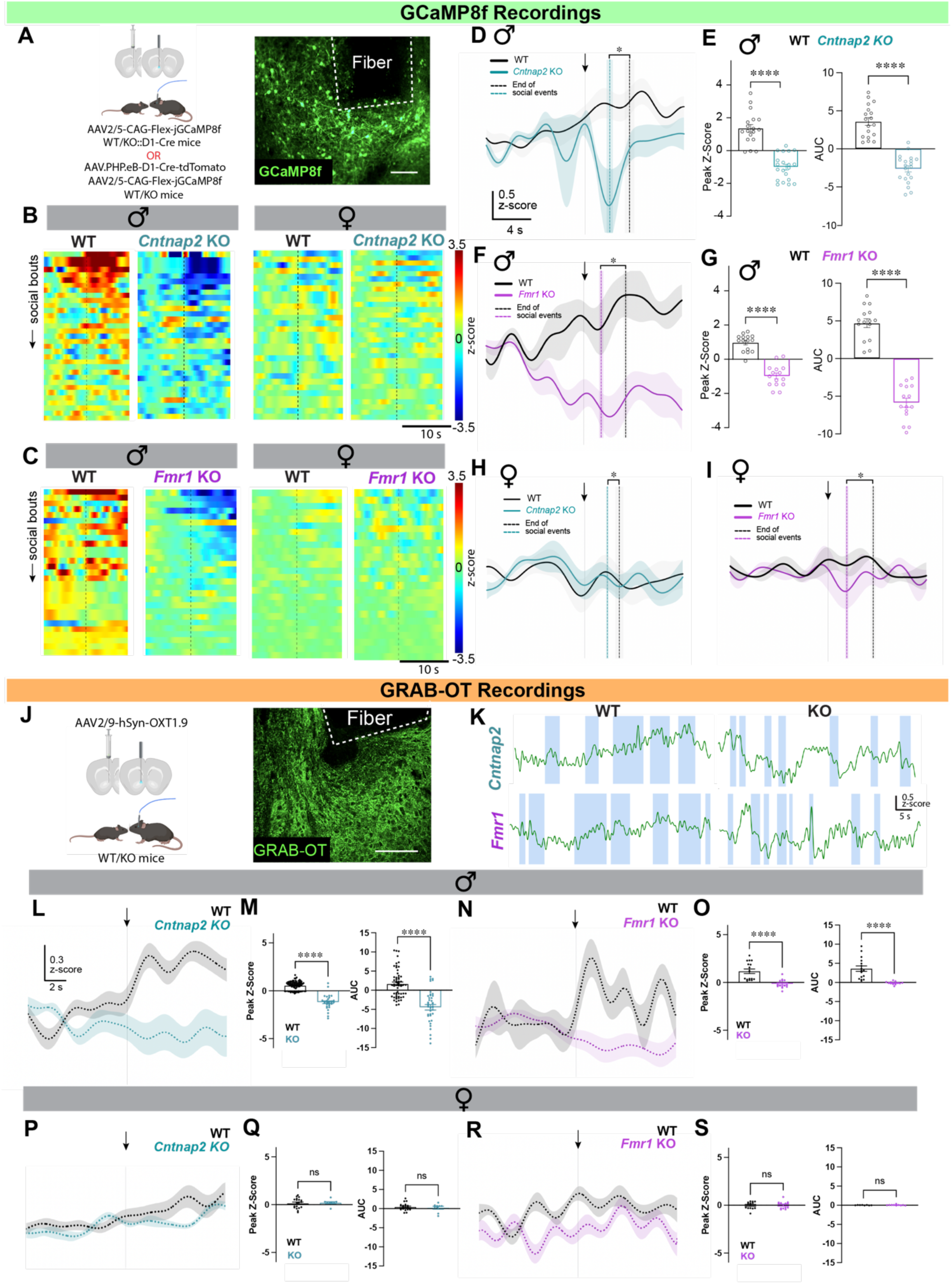
*Cntnap2* KO and *Fmr1* KO males exhibit reduced NAcSh D1-MSN activity and a lack of OXT release during social investigation. **a.** Schematic: Viral strategy to target D1-MSNs (left), representative micrograph showing GCaMP8f expression and fiber implant targeting (right, scale bar=100μm). **b-c.** Sex- and genotype-stratified GCaMP8f signal heatmaps during social investigation. Dotted line in the center represents social onset. **d,f,h,i**. Averaged fiber photometry traces (+/- SEM shaded) 10 seconds before and after onset of social investigation (t=0s, arrow) showing male-specific genotype differences in Ca^2+^ dynamics during social investigation (*Cntnap2*: green, *Fmr1*: purple, WT: black). Vertical dotted lines represent average end-times for social bouts (+/- SEM shaded). **e,g.** Comparisons of AUC and peak z-scores between WT and KO males. Two-sample unpaired t-test. ****P<0.0001. **j.** Schematic: Viral strategy to express GRAB-OT (*left*), representative micrograph showing GRAB-OT expression and fiber implant targeting in the NAcSh (*right*, scale bar=100μm). **k.** Example traces showing GRAB-OT signals during social investigation (bouts shaded in blue). **l,n,p,r.** Averaged fiber photometry traces (+/- SEM shaded) 10 seconds before and after social investigation onset (t=0s, arrow) showing sex and genotype differences in OT dynamics in the NAcSh during social investigation (*Cntnap2*: green, *Fmr1*: purple, WT: black). **m,o,q,s.** Sex stratified peak z-scores and AUC. Two-sample t-tests. *P<0.05, **P< 0.01, ****P<0.0001, ns=not significant.

The NAcSh receives dense projections from PVN-OXT neurons ^63,64^. We and others have demonstrated that both OXT injection into the NAcSh^68–70^, and optogenetic activation of NAc-projecting PVN-OXT fibers can drive increased NAcSh activity and modify social behavior^19^. To directly determine whether social investigation elicits OXT release in the NAcSh, we measured OXT release in the NAcSh using GRAB-OT, a GPCR-based genetically encoded fluorescent OXT sensor^71^, which we and others have previously used to detect OXT release dynamics during social investigation ^71,72^. In line with our above GCaMP8f data (**Fig.4**), we found a striking sex difference in OXT release dynamics in the NAcSh during social investigations, wherein WT males showed significantly larger social investigation-induced changes in GRAB-OT signal than females (**Fig.4l-s**). In contrast to WT males, both *Cntnap2^-/-^* and *Fmr1^-/Y^* males showed GRAB-OT signals that decayed below baseline close to social investigation onset (p*_Cntnap2_* < 0.0001, p*_Fmr1_* <0.0001, Two sample unpaired t-test; **Fig.4l-o, Table.4**) and showed no change in AUC values (Two sample unpaired t-test; **Fig.4m,q**).

### Cntnap2 KO and Fmr1 KO females exhibit reduced GABAergic neuronal activity and lack social investigation-induced OXT release in the CeL

Our unexpected observation of that the NAcSh D1-MSNs did not encode social stimuli during female-female interactions suggested a separate female-specific circuit mechanism, where the activity level of another brain region regulates the length of social engagement. The amygdala is another SSN region receiving dense projections from PVN-OXT neurons^73^, and encodes the salience of social stimuli^74^. In particular, OXT modulation in the CeA has female-specific effects on social behavior^75,76^, and CeA activity level is strongly predictive of the duration of social investigation bouts in both KO models as we have shown above (**Fig.5**). Of the two subdivisions within the CeA, the CeL receives heavier excitatory input from PVN-OXT neurons compared to its medial counterpart^73^ and inhibiting CeL GABAergic neurons reduces the duration of social interaction in WT mice^62^. Therefore, we hypothesized that GABAergic CeL neurons in KO mice exhibit aberrant activity during social investigation in a female-specific manner. To test this, we measured population-level GCaMP8f calcium dynamics in CeL GABAergic neurons via fiber photometry in KO mice and their WT littermates during social investigation.

**Figure 5.**
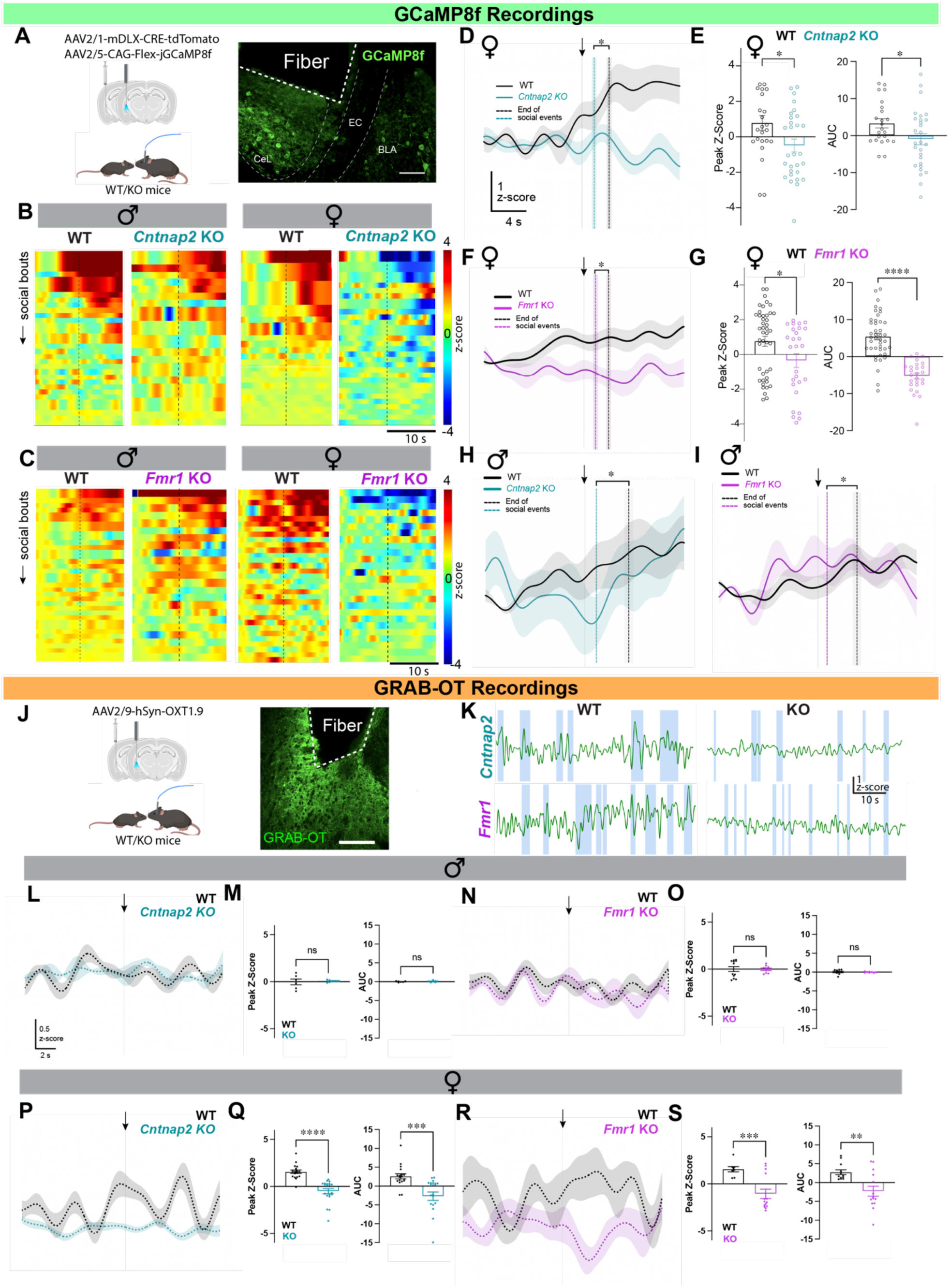
*Cntnap2* KO and *Fmr1* KO females exhibit reduced CeL GABAergic neuronal activity and lack of OXT release during social investigation. a. Schematic: Viral strategy to target GABAergic neurons (left), representative micrograph showing GCaMP8f expression and fiber implant targeting (right), EC, external capsule. BLA, basolateral amygdala. Scale bar=100μm. **b-c.** Sex- and genotype-stratified GCaMP8f signal heatmaps during social investigation. Dotted line in the center represents social onset. **d,f,h,i.** Averaged fiber photometry traces (+/- SEM shaded) 10 seconds before and after onset of social investigation (t=0s, arrow) showing female-specific genotype differences in Ca^2+^ dynamics during social investigation (*Cntnap2*: green, *Fmr1*: purple, WT: black). Vertical dotted lines represent average end-times for social bouts (+/- SEM shaded). **e,g.** Comparisons of AUC and peak z-scores between WT and KO males using Two-sample unpaired t-tests. ****P<0.0001. **j.** Schematic: Viral strategy to express GRAB-OT (*left*), and representative micrograph showing GRAB-OT expression and fiber implant targeting in the CeL (*right*, scale bar=100μm.). **k.** Example traces showing GRAB-OT signal. **l,n,p,r.** Averaged fiber photometry traces (+/- SEM shaded) 10 seconds before and after social investigation onset (t=0s, arrow) showing sex and genotype differences in OT dynamics in the CeL during social investigation (*Cntnap2*: green, *Fmr1*: purple, WT: black). **m,o,q,s.** Sex stratified AUC and peak z-scores analysis of WT and KO mice in the CeL using two-sample t-tests. *P<0.05, **P<0.01, ****P<0.0001. ns=not significant.

We found a gradual increase in GCaMP8f signal from CeL GABAergic neurons prior to the onset of social investigation in both male and female WT mice (**Fig.5b-d,f**). In alignment with our above c-Fos data (**Fig.3**), this canonical increase in social-related activity was intact in both *Cntnap2^-/-^*and *Fmr1^-/Y^* males (**SFig.3**), despite these mice having shorter social bout lengths compared to their WT littermates (p*_Cntnap2_*<0.0001, p*_Fmr1_*<0.0001, Two-sample unpaired t-test, **SFig.5**). Indeed, there were no sigificant differences in peak values and AUCs in male and female WT mice as well as male KO mice (**Fig.5e,g, Table.5**). However, in both *Cntnap2^-/-^*and *Fmr1^-/-^* females, GCaMP8f signal remained at baseline and no peaks were detected (**Fig.5, Table.5**). Alongside this lack of CeL engagement, KO females also showed shorter individual social bouts than their WT littermates (unpaired t-test, p*_Cntnap2_*=0.0003, p*_Fmr1_*=0.0006, **SFig.5**).

Given that OXT release dynamics in the NAcSh of *Cntnap2^-/-^*and *Fmr1^-/-,-/Y^* mice are selectively affected in males and align well with the sex-specific neuronal activity phenotypes, we hypothesized that OXT-release dynamics in the CeL of *Cntnap2^-/-^* and *Fmr1^-/-,-/Y^* mice were affected in a female-specific manner. Indeed, we found a sex difference in OXT release dynamics in the CeL during social investigation. *Cntnap2^+/+^* and *Fmr1^+/+^* females showed significantly larger social investigation-induced changes in GRAB-OT signal compared to *Cntnap2^-/-^* females, which showed minimal signal changes from baseline, and *Fmr1^-/-^* females, which showed a decrease in GRAB-OT signal below baseline close to the onset of social investigation (unpaired t-tests, p_Peak*Cntnap2*_ < 0.0001, p_Peak*Fmr1*_ =0.0008, p_AUC*Cntnap2*_ = 0.0016, p_AUC*Fmr1*_ =0.0012; **Fig.5l,m,p,q, Table.5**). Meanwhile, WT and KO males across genotypes showed a similar lack of deviation from baseline in GRAB-OT signal (p_Peak*Cntnap2*_=0.7999, p_Peak*Fmr1*_ =0.9739, p_AUC*Cntnap2*_ =0.6863, p_AUC*Fmr1*_ =0.7386; **Fig.5n,o,r,s, Table.5**).

In summary, both WT and KO males show increased CeL activity during social investigation. WT females also show this canonical circuit engagement, whereas this response was absent in KO females. Furthermore, social investigation induced OXT release in the CeL only in WT females, a response also absent in KO females. Together, these data suggest that deficient OXT recruitment of CeL activity may contribute to the shorter social investigation bouts observed in female KO mice.

### Restoring OXT signaling in the NAcSh and CeL is sufficient to rescue activity and social behavior phenotypes in male and female KO mice respectively

Having observed insufficient OXT release during social investigation in KO mice in a sex- and region-specific manner that is also consistent with circuit activity aberrations, we predicted that evoking OXT release (via optogenetic stimulation of OXT fibers) in a region-specific manner would rescue both social behavior and circuit activity phenotypes in a sex-dependent manner. To test this, we expressed ChrimsonR^77^, a red-light activated channelrhodopsin, in PVN OXT neurons, then light-stimulated OXT fibers in the NAcSh or CeL while also performing GCaMP8f recordings during home cage social investigation. Indeed, we found that 5-minute-long, unilateral light stimulation of ChrimsonR-expressing OXT fibers in the NAcSh successfully rescued the aberrant activity in D1-MSNs in both *Cntnap2^-/-^*and *Fmr1^-/Y^* males. In the 5-minute period following optogenetic stimulation, these mice showed increased GCaMP8f signal close to social investigation onset (**Fig.6d**), positive peak z-scores and positive AUC values compared to prior baseline GCaMP8f signals from the same mice without optogenetic stimulation (One-sample Welch’s t-test; **Fig. 6d, Table.6a**). Simultaneously with the GCaMP8f signal changes, the mean duration of social bouts in both *Cntnap2^-/-^*males and *Fmr1^-/Y^* (males) also increased (q*_Cntnap2_* = 0.006, q*_Fmr1_* = 0.0022; Two-way repeated-measures ANOVA, FDR correction; **Fig.6f, Table.6a**). In contrast, optogenetic stimulation of OXT fibers in the NAcSh had no effect on D1-MSN activity or the duration of social investigation bouts in either *Cntnap2^-/-^* females or *Fmr1^-/-^* females (**Fig**.**6e**).

**Figure 6.**
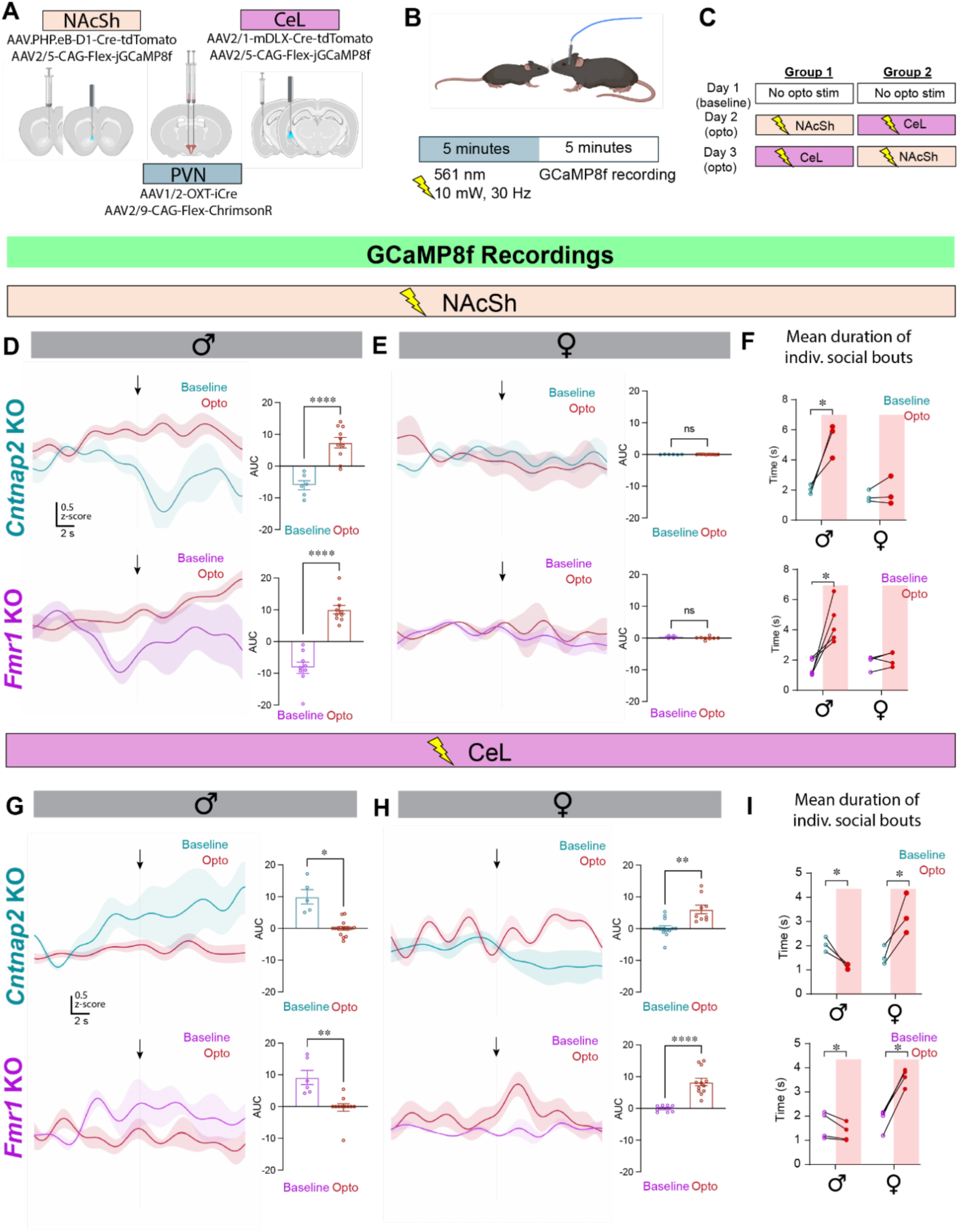
Restoring OXT signaling in the NAcSh and CeL is sufficient to rescue neuronal activity and social behavior phenotypes in male and female KO mice respectively. **a.** Schematic: Viral strategy for optogenetic stimulation of OT fibers in the NAcSh or CeL. **b, c.** Within-subject experimental design for GCaMP8f recordings from NAcSh and CeL after optogenetic stimulation. **d,e,g,h.** Averaged fiber photometry traces (+/- SEM shaded) 10 seconds before and after onset of social investigation (t=0s, arrow) showing a male-specific rescue of social-induced neuronal activity in the NAcSh (**d,e**) and a female-specific rescue of social-induced neuronal activity in the CeL (**g,h**) of KO mice (*Cntnap2* KO baseline: green, *Fmr1* KO baseline: purple, activity after optogenetic stimulation in NAcSh, CeL: red). Two-sample t-tests, *P<0.05, **P<0.01, ****P<0.0001. ns=not significant. **f,i.** Sex-stratified comparisons (across treatment conditions) of mean duration of social bouts. Two-way repeated measures ANOVA with FDR correction for multiple comparisons, *q<0.05.

Conversely, we found that after 5 minutes of unilaterally stimulating OXT fibers in the CeL, GABAergic neurons in both *Cntnap2^-/-^*and *Fmr1^-/-^* females showed increased GCaMP8f signal close to social onset compared to no-stimulation baseline signals from the same mice (One-sample t-tests; **Fig.6g,h, Table.6b**). 5 minutes of OXT fiber stimulation in the CeL also significantly increased the mean duration of social bouts in both *Cntnap2^-/-^* and *Fmr1^-/-^* females (p*_Cntnap2_* = 0.0136, p*_Fmr1_* < 0.0001, Two-way repeated measures ANOVA with FDR correction; **Fig**.**6i, Table.6b**). Interestingly, in *Cntnap2^-/-^* and *Fmr1^-/Y^* males, optogenetic stimulation of OXT fibers in the CeL prevented the expected increase in GABAergic neuronal activity (peak z-scores and AUC values close to zero; one-sample Welch’s t-test; **Fig.6h**), and further reduced social investigation bout duration in these KO males (p*_Cntnap2_* = 0.0465, p*_Fmr1_* = 0.041; two-way repeated measures ANOVA with FDR correction; **Fig.6i, Table.6b**).

Collectively, these findings demonstrate that distinct ASD-risk gene mutations converge on deficient OXT recruitment of sexually dimorphic social circuits, with restoration of OXT signaling in the NAcSh of males and CeL of females rescuing both circuit activity and social engagement.

## Discussion

Using chemogenetics, c-Fos immunolabeling, SHIELD whole-brain clearing, and optogenetics, we demonstrate a network- and region-level convergence in aberrant SSN activity between two independent KO models of ASD-risk gene disruptions: *Cntnap2^-/-^* and *Fmr1^-/-,-/Y^*mice. Furthermore, we show that endogenous OXT release alters SSN network connectivity and regional activity phenotypes to consequently rescue disrupted social behavior in a model convergent manner. Population-level activity dynamics during social investigation revealed convergent behavioral and circuit-level phenotypes that are mediated by sex-specific neural substrates, namely aberrant activity in the NAcSh in males and the CeL in females. Further, we demonstrate that this aberrant activity is accompanied by insufficient OXT release in affected regions, and that optogenetically evoked OXT release in those regions rescues both circuit activity and social behavior phenotypes in KO mice.

### Convergent behavioral phenotypes in maintaining social investigation bouts

In our study, we performed a systematic evaluation of the social investigation behavior of male and female *Cntnap2^-/-^* and *Fmr1^-/-,-/Y^* mice while they were freely interacting with sex-matched conspecifics and compared them with their WT littermates. We found that reduced sociability phenotypes in both *Cntnap2^-/-^* and *Fmr1^-/-,-/Y^* mice are driven by a lack of interest in continuing social investigation initiated at a normal number (**Fig.1**). These consistent social phenotypes across the two models and sexes are notable, given that recent studies in *Cntnap2^-/-^* and *Fmr1^-/-,-/Y^* mice have yielded inconsistent reports on sex differences in social phenotypes, with some studies showing no disruption in social behaviors in KO females^28,36^, while others report hypersociability, particularly in *Fmr1^-/-^* females^78^. We resolved these inconsistencies by demonstrating that compared to *Fmr1^+/+^* females, *Fmr1^-/-^*females show a high number of very short social investigation bouts, obscuring their social phenotype when measured simply as the total duration of social investigation over a 10-minute period (**Fig.1**), thus highlighting the sensitivity of our experimental approach in identifying shared disruptions in social phenotypes across models and sexes. Furthermore, the WT littermates for each KO model exhibited extremely consistent social behavior in our dataset (**Fig.1**), which increases confidence in the shared aspects of the social behavioral phenotypes of the two KO models.

### An SSN subnetwork crucial for encoding the duration of social engagement

Our data show a strikingly strong relationship between the length of social investigation bouts and cellular activity in several SSN regions, underscoring the functional role of these regions in the maintenance of social engagement. Specifically, the cellular activity levels in 4 key SSN regions (NAc, CeA, ILA, PVN), assessed as the number of c-Fos+ cells, were positively correlated with the duration of social investigation bouts in both KO models. These results are consistent with the proposed role of these SSN nodes in facilitating sustained social interactions^50,62,79,80^. Previous studies have shown that the ILA has a descending excitatory influence on the activity of the CeA and the NAc^81,82^, which have been shown to facilitate social interaction^83^. Additionally, the ILA, CeA, and NAc all receive fiber projections from OXT neurons in the PVN, which are activated by social stimuli and potentiate their neuropeptide release onto these downstream targets^61,84^. Therefore, it is likely that the coordinated activity of the ILA-NAc and ILA-CeA circuits contributes to the rewarding nature of high-quality social interactions in WT mice, further potentiated by the neuromodulatory effects of OXT released by PVN-OXT neurons. In contrast, social investigation failed to activate these 4 SSN regions in both KO models and the functional connectivity across these regions remains weak (**Fig.2,3**), but these phenotypes are strikingly rescued by chemogenetic stimulation of endogenous OXT release (**Fig.2**). Therefore, these data underscore the increased activity and functional connectivity of this SSN subnetwork as a key mechanism underlying OXT-mediated behavioral rescue in both KO models.

### OXT release dynamics in NAcSh and CeL during social investigation

We report striking sex differences in regional OXT release during social investigation: male-specific in the NAcSh and female-specific in the CeL (**Fig.5**). The effects of exogenously administered OXT in the NAcSh and CeL are sexually dimorphic^85,86^, which could indicate differential OXT release *in vivo*. A rodent fMRI study reported that intracerebroventricular administration of OXT caused more activation in the NAcSh in males and more activation in the CeA in females^85^, which aligns well with our findings. OXT also enhances GABAergic transmission in CeL, with females showing higher CeL OXT receptor mRNA expression than males^86^. Interestingly, optogenetic activation of OXT fibers (leading to putative OXT release) in the CeL concomitantly increased the activity of its GABAergic neurons and the duration of social investigation in female mice, while the same manipulation in male mice suppressed the canonical WT-like CeL activity increase during social investigation and also shortened the duration of social investigation, exacerbating their social phenotype (**Fig.4,6**). The unexpected antagonistic relationship between OXT signalling and social engagement that we observed exclusively in male CeL is consistent with a previous report that identified a negative correlation between OXT release (measured via microdialysis) in the CeA and the duration of social investigation in males^60^. Because the CeL shows intact activation in male KO mice during social investigation (**Fig.4**) with an absence of OXT release similar to WT males (**Fig.5**), perhaps additional optogenetically evoked OXT release in this region has a counterproductive effect on CeL activity. In summary, these results highlight that deficient OXT release in KO models may be restricted to specific circuits depending on the sex, and despite other evidence of a dysfunctional central OXT system in these mice, the bidirectional control of social engagement appears to be preserved at least in the CeL.

### NAcSh and CeL encoding of social stimuli and sex differences in WT and KO mice

We report, for the first time, a striking male specificity in the recruitment of NAcSh D1-MSNs in WT mice engaging in social investigation in a home cage setting (**Fig.4**). This observation can potentially be attributed to the NAc’s role in modulating dominance behaviors during social interactions^79,87,88^, which are more pronounced in males^88–90^, compared to females^88^. Although we did not observe any changes in NAcSh activity during female-female interactions, more intense social competition^88,91^ could result in NAcSh recruitment in both sexes. Interestingly, both *Cntnap2^-/-^* and *Fmr1^-/Y^* males experience consistent social subservience, observed as more frequent social defeats in *Fmr1^-/Y^* males^92,93^ and decreased territorial behaviors in *Cntnap2^-/-^* males^94^. This social “defeat” could thus be associated with inhibition of D1-MSNs in the NAcSh (**Fig.4**), while WT males, who are more likely to “win” during homecage interactions, exhibit the opposite activity pattern. In contrast to the NAc, we observed that CeL GABAergic neurons were activated in WT mice of both sexes (**Fig.5**), consistent with previous findings in male mice, in which manipulations of CeA activity bidirectionally modulated social contact duration^62,95^. While CeA activity levels in socially interacting females have not been reported, our results are consistent with previous studies showing that experimentally increasing CeA activity facilitates socioemotional discrimination^73^ and reward processing^62^ in both sexes. Interestingly, social investigation appears to inhibit CeL activity only in female *Cntnap2^-/-^*and *Fmr1^-/-^* mice (**Fig. 5**). The preservation of increased CeL activity in male KO mice, despite their behavioral phenotype of shorter social investigation bouts, reveals that the CeL-dependent component of social investigation behavior may be intact in males. Indeed, the NAc and the CeL have been suggested to encode distinct components of reward processing, where the CeA drives “incentive motivation” while the NAc encodes the “hedonic payout” component of reward (as reviewed in ^96, 65^). Given this framework, our results suggest that CeL-incentive encoding is intact while the NAc-dependent hedonic payout is impaired in KO males, leading to shorter social bouts. This possibility is further supported by our results in **Fig.6**, where optogenetic stimulation of OXT release in the CeL suppressed CeL activity and further exacerbated the social phenotype in males. Collectively, these data imply a potential sex bias in which component(s) of social reward are affected in *Cntnap2^-/-^* and *Fmr1^-/-,-/Y^* mice, where a disrupted maintenance of social investigation may be associated with social “anhedonia” in males and social “indifference” (no/low valence) in females.

### Concluding remarks

Our comprehensive analysis of circuit pathologies observed in gene-KO models of two functionally distinct ASD-risk genes has revealed striking model-convergent sex differences in: 1. SSN regions recruited during social behavior in genetically intact mice, 2. region-specific disruptions in neuronal activity during social investigation, 3. region-specific OXT release dynamics during social investigation, 4. region-specific circuit modulation and behavioral rescue via evoked local OXT release. We provide compelling evidence that sex-dependent OXTergic modulation of SSN circuits is implicated in facilitating sustained social investigation and that restoring OXT signaling in affected circuits is sufficient to rescue both circuit and behavioral phenotypes caused by ASD-linked genetic mutations. Collectively, these findings identify sex-dependent OXTergic modulation as a likely unifying circuit mechanism linking genetic risk to social dysfunction in ASD cases that feature central OXT system dysfunction.

## Methods

### Animals

All animal protocols were approved by the McMaster University Animal Research Ethics Board (AREB). Drd1-cre mice (B6;129-Tg(Drd1-cre)120Mxu/Mmjax, RRID:MMRRC_037156-JAX)^97^ were obtained as initial breeders from the Mutant Mouse Resource and Research Center (MMRRC) at The Jackson Laboratory, an NIH-funded strain repository, and was donated to the MMRRC by Ming Xu (The University of Chicago). *Cntnap2* KO mice (B6.129(Cg)- *Cntnap2^tm1Pele^*/J (stock #017482))^98^ was obtained as initial breeders from Jackson Laboratories, and was donated by Elior Peles (Weizmann Institute of Science). *Fmr1* KO mice (B6.129P2-*Fmr1^tm1Cgr^*/J (stock #003024)^99^ was obtained as initial breeders from Jackson Laboratories, and was donated by Ben Oostra (Erasmus University) and Stephen Warren (Emory University School of Medicine). These mice were then bred in-house in a trio (1 male: 2 females) as follows.

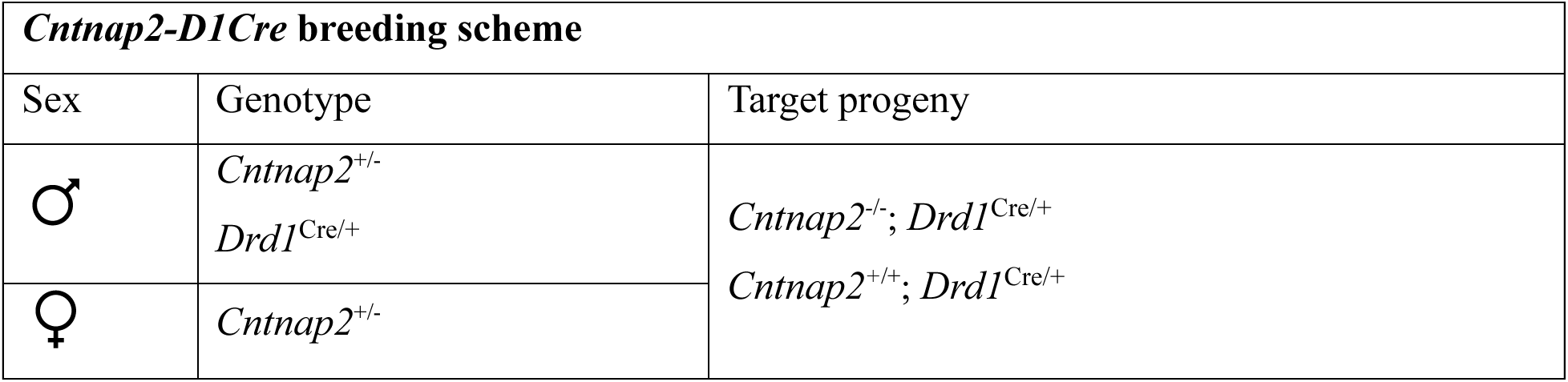

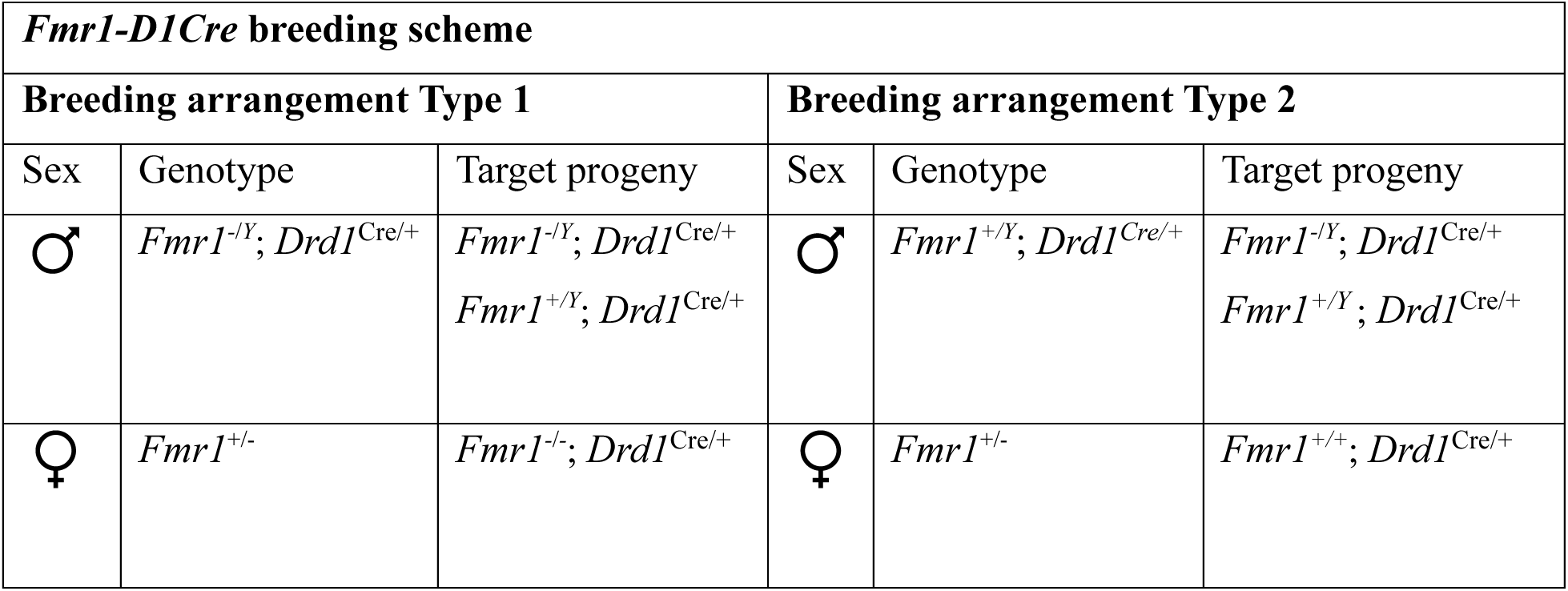

The genotype for each mouse was determined using PCR analyses as described by Jackson Labs for each strain. We used the following primers:

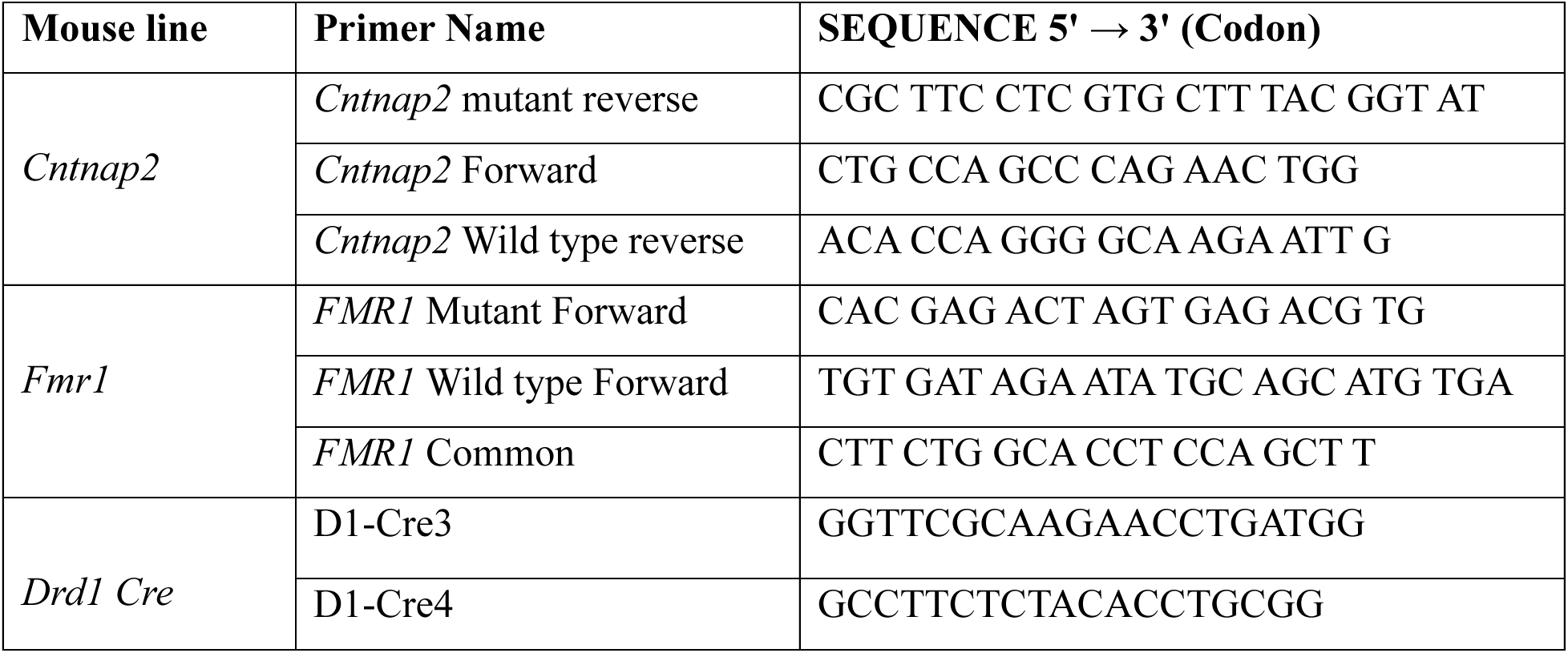

Mice were housed in polycarbonate cages (28cm x 17.5cm x 12cm) with bedding, nesting material, and an enrichment object (plastic tube). Breeders were provided with extra nesting material. Mice were housed with 1-3 sex matched and strain matched littermates. Food and water were available *ad libitum*. The mouse vivarium was kept at a 12-hour day/night cycle (lights on from 8AM-8PM) and an ambient temperature of ∼22°C.

Stereotaxic virus injection surgeries were conducted when mice were 8-13 weeks old. Behavior experiments were conducted when mice were 10-16 weeks old. Stimulus mice were sex-matched WT juveniles that were novel to experimental mice (4-5 weeks old, bred from WT x WT pairings separately from KOs).

### Behavioral experiments

#### Handling and habituation

All behavior testing was conducted by the same experimenter. Mice were handled by the experimenter for 3 consecutive days preceding behavior testing, for 3-5 minutes on each day, to habituate the mice to the experimenter. On the day of behavior testing, mice were socially isolated (to enhance social motivation) in a home-cage environment in the testing room for 1 hour (to allow habituation to the testing room). Food and water were available *ad libitum*. During this time, the behavior testing room was maintained at dim light (∼2 lumens), with a white noise machine (∼70 dB) and an air purifier to filter strong odors and dander (Honeywell). These room conditions were maintained throughout the social behavior test.

#### Home cage social investigation

To assess social investigation differences between KO mice and their WT littermates, we performed the home cage social investigation test similar to previously published protocols^19,28^ using a custom-built three-chambered apparatus with opaque walls (60cm x 40.5cm x 22cm, 20cm inner chamber width). Each chamber contained an open standard home cage with bedding only (no nesting material or enrichment). Behavior was recorded using an overhead camera (Chameleon3 CM3-U3-31S4M; Sony IMX265, 1/1.8” Mono CMOS sensor) for offline analysis with the event capture software BORIS^100^.To assess social investigation differences between KO mice and their WT littermates, we performed the home cage social investigation test similar to previously published protocols^19,28^ using a custom-built three-chambered apparatus with opaque walls (60cm x 40.5cm x 22cm, 20cm inner chamber width). Each chamber contained an open standard home cage with bedding only (no nesting material or enrichment). Behavior was recorded using an overhead camera (Chameleon3 CM3-U3-31S4M; Sony IMX265, 1/1.8” Mono CMOS sensor) at a frame rate of 17 frames/s.

First, mice went through a 10 min habituation phase where the experimental mouse and the juvenile stimulus mouse were placed in separate home cage with fresh bedding. This was followed by a 10 min social investigation test phase where the experimental and stimulus mouse were placed together in a new homecage with fresh bedding, and allowed to interact freely for the duration of the phase.

For WT-KO comparisons of social investigation-elicited cellular activity in the CeA, NAc, PVN, and ILA, a group of mice were subjected to a 10-minute home cage social investigation as described above (“home cage + social investigation” group). Another group of mice were placed alone in a new home cage setting without any stimulus mouse (“home cage” group) to serve as controls.

### Stereotaxic Surgeries

Stereotaxic surgeries were performed to intracranially deliver AAVs and/or implant optic fibers using previously described protocols^19^. Mice were head-fixed on a stereotaxic apparatus and maintained under isoflurane gas anaesthesia (∼1-2%) delivered continuously via a nose cone (approx. surgery duration was 1.5-2 hours). Post-surgery, mice were single housed overnight with extra enrichment (wooden tongue depressor) post-surgery to reduce risk of re-opening of the incision site by littermate overgrooming. Mice were also treated with carprofen (5mg/kg) for 3 consecutive days following surgery for pain relief. All behavior tests were conducted 3-4 weeks following surgery.

For intracranial injections, AAVs were infused at a rate of 0.02µL per minute using a 2µl Neuros syringe (Hamilton Company). The virus was allowed to diffuse for 5 minutes before the needle was gradually withdrawn.

For chemogenetic stimulation of endogenous OXT release, mice were injected with either AAV2-mOXT-Venus^101^ (plasmid generation by the Grinevich laboratory, then viral production by Signagen) or AAV2*-*OXT-hM3Dq-mCherry^28^ (plasmid generation by the Geschwind laboratory, UCLA, then viral production by Signagen) was bilaterally injected into the PVN [0.3µl/side, AP: −0.6, ML: +/-0.2, DV: −4.8].

For NAcSh photometry cohorts, GCaMP8f expression was targeted to D1-expressing MSNs via stereotaxic injection of AAV2/5-CAG-FLEX-jGCaMP8f (COVF viral vector core, RRID:SCR_016477)^102^ into *Cntnap2*^-/-^; *Drd1*^Cre/+^ or *Fmr1*^-/-.-/Y^; *Drd1*^Cre/+^ mice or by a combined injection of two AAVs (1:1) each carrying a D1-Cre recombinase construct and a Cre-dependent GECI (AiP13779 -AAV.PHP.eB-AiE0779m_3xC2-minBG)-iCre(R297T)-BGHpA^103^ (Addgene) + AAV2/5-CAG-FLEX-jGCaMP8f) into *Cntnap2*^-/-^ or *Fmr1*^-/-,-/Y^ mice. For CeA photometry cohorts, GCaMP8f expression was targeted to GABAergic neurons via a combined injection of two AAVs (1:1) each carrying a mDlx-Cre recombinase construct and a Cre-dependent GECI (AAV2/1-mDLX-CRE-tdTomato (COVF Viral Vector Core) + AAV2/5-CAG-FLEX-jGCaMP8f). Because the CeL in mice is lateralized (as reviewed in^104,105^), and the CeA in the right hemisphere is associated with pain modulation^106^, we targeted the left CeL, which has been shown to modulate social behavior^107^. For the purpose of consistency, we also targeted the left NAcSh. The AAV was injected unilaterally (0.3µl in the left hemisphere) targeting the medial shell of the NAc (AP: +1.18mm; ML: −0.5mm; DV: −4.0mm) or the CeL (AP: −1.46, ML: −3.05, DV: −4.51).

For optogenetic stimulation of OXT neurons, an AAV carrying an OXT-Cre construct (AAV1/2-OXT-iCre^108^, produced in house at the Grinevich laboratory) and another AAV carrying a construct for Cre-dependent expression of ChrimsonR^77^, a red-shifted excitatory opsin (AAV2/9-CAG-Flex-frwd-ChrimsonR-GSG-P2A-tdTomato, COVF viral vector core), were mixed 1:1 and injected bilaterally into the PVN (0.3µl on each side, AP: −0.6, ML: +/-0.2, DV: −4.8)^77^, a red-shifted excitatory opsin (AAV2/9-CAG-Flex-frwd-ChrimsonR-GSG-P2A-tdTomato, COVF viral vector core), were mixed 1:1 and injected bilaterally into the PVN (0.3µl on each side, AP: −0.6, ML: +/-0.2, DV: −4.8).

For OXT sensor recordings, 0.3µl of AAV2/9-hSyn-OXT1.9 (GRAB-OT, plasmid generation by Yulong Li laboratory, Peking University, viral production by BrainVTA)^109,110^ was injected unilaterally into the left NAcSh (AP: +1.18mm; ML: −0.5mm; DV: −4.0mm) and/or the left CeL (AP: −1.46, ML: −3.05, DV: −4.51).

For all fiber photometry cohorts, following virus injection, a single optic fiber was implanted to target either the left NAcSh and/or left CeL for single-site GCaMP8f/GRAB-OT recordings. For optogenetic stimulation cohorts, two optic fibers, each targeting either the NAcSh or CeL for dual-site recordings, were installed at the same coordinates as virus injection. The implant was affixed to the skull using Metabond (Parkell). After the Metabond had completely hardened, the incision site was closed around the Metabond with Vetbond skin adhesive.

### Chemogenetic activation of PVN-OXT neurons

To avoid unspecific c-Fos-activation associated with the potential stress of intraperitoneal (*i.p.*) injection of clozapine-n-oxide (CNO, Tocris) administered prior to behavioral testing, we used a two-step protocol: the day prior to behavioral experimentation, mice were given a single dose (1mg/kg) of *i.p.* CNO. Then, they were single housed overnight, in which they had free access to CNO in drinking water (8 mL of water containing 4 µg CNO/ml), contained in a small glass tube with rubber stopper spouts (AMUZA Drinko measuring sipper tubes) to be able to measure the exact amount of daily water consumption by each mouse. The tube was wrapped in foil to protect against the degradation of CNO which is light sensitive. This protocol was adapted from a previously published protocol^111^ to maintain a dose of 1mg/kg overnight through ∼5 mL of water intake, which is the typical daily amount consumed by an adult male mouse^112^. Drinko tubes were weighed before and after experimentation to calculate CNO water consumption.

### Whole brain c-Fos immunolabeling using SHIELD tissue clearing

∼100 minutes following home cage social investigation, mice were transcardially perfused with 4% paraformaldehyde in 1X phosphate buffer (Fujifilm Biosciences) for further histological analysis. This time point was chosen based on previously published protocols ^113,114^ for measuring peak c-Fos expression. The brains were then processed for SHIELD tissue clearing and c-Fos immunostaining. SHIELD preservation and active de-lipidation were conducted following our previously published whole-brain IHC protocols^115^. Following delipidation, brains were blocked in 5% NDS in Antibody Blocking Solution (LifeCanvas ABS-250) at 37°C with shaking for 2 days. We processed 10 brains in every SHIELD processing cohort.

For active primary immunolabeling, samples were pre-incubated in SmartBatch+ Radiant Buffer (LifeCanvas #SB-RB-500) for 3 days with gentle shaking at room temperature to enhance probe penetration, refreshing the buffer on the morning of labeling and continuing incubation for at least 1 h prior to setup. Staining cups were leak-tested, rinsed with tap and distilled water, and filled with Radiant Buffer (40 mL in a large batch cup) to which we added anti-c-Fos (24µg, c-Fos-Rabbit, Abcam EPR20769), anti-GFP (40µg, GFP-Chicken, Aves), and anti-RFP (40µg, RFP-Goat, Rockland) primary antibodies. Samples were placed in mesh bags, submerged in the buffer, and loaded into the SmartBatch+ device equipped with the

Radiant pH Catalyst (LifeCanvas #SB-RC) and column. Electrophoretic labeling was performed at 30°C under preset conditions (90 V, 350 mA) for 35 hours with continuous rotation. On completion, samples were transferred to PBS with 0.02% sodium azide (PBSN) for washing (2x for ∼3 hours), then fixed in 4% paraformaldehyde at room temperature overnight.

Secondary labeling with anti-rabbit (48µg Alexafluor+ 647, Jackson Immuno), anti-chicken (80µg Alexafluor+ 488, Jackson Immuno), and anti-goat (80µg Alexafluor+ 555, Jackson Immuno) secondary antibodies and index matching was performed using the same general protocol as published before^115^.

### Whole brain Lightsheet imaging

Index-matched samples were embedded in an index-matched agarose block and placed in an imaging chamber filled with immersion oil imaging medium (LifeCanvas #E.1). We collected whole brain images using a SmartSPIM axially swept lightsheet microscope by LifeCanvas Technologies. We used 647 nm, 488 nm and 561 nm excitation lasers with a 2 μm Z-interval, employing a 3.6× objective lens (resulting voxel size: 1.8 μm × 1.8 μm × 2.0 μm). Individual tiles were corrected for striping artifacts using DestripeGUI (LifeCanvas Technologies), followed by image stitching in StitchGUI (LifeCanvas Technologies). The stitched datasets were subsequently assembled into image stacks with Microfile+ (MBF Bioscience) for subsequent analysis.

### Tissue section c-Fos immunolabeling and fluorescence imaging

Two groups of mice, “home cage + social investigation” and “home cage” (behavioral task described above) were transcardially perfused with 4% paraformaldehyde (PFA) in phosphate buffer saline (PBS) ∼100 minutes after behavior. PFA-fixed brains were extracted and serially sectioned on a vibratome into 50µm thick sections. Representative sections from each SSN ROI (NAc, PVN, CeA, ILA) were immunostained to label c-Fos immunopositive (c-Fos+) cells (Primary antibody 1:200: xc-Fos-Rb, Synaptic Systems; Secondary 1:500: AlFl+ xRb-555, ThermoFisher). Labeled sections were mounted with Fluoromount-G mounting medium with DAPI (Invitrogen) for nuclear staining and imaged at 10× and 0.75 pixels/μm using Zeiss Axioscan 7 slide scanner at the McMaster Centre for Advanced Light Microscopy (CALM).

### Optogenetics and Fiber Photometry Recordings

#### Fiber photometry recordings of GCaMP8f and GRAB-OT signals

Fiber photometry recordings of GCaMP8f and GRAB-OT1.9 signals in the NAcSh and CeL were performed in separate cohorts during home cage social investigation as above, with the following modifications. During the habituation period the experimental mouse’s fiber implant was coupled with a pliant overhead fiber-optic patch cable, allowing for the mouse to habituate to the patch cable for 10 minutes. No lasers or LEDs were on, and no imaging was conducted during the habituation phase. Fiber photometry recording began just before the experimental and stimulus mouse were placed in the center cage for the testing phase. GCaMP8f or GRAB-OT recordings were performed during the 10-minute testing phase using a Doric fiber photometry system with two light-emitting diodes (LEDs; isosbestic, 405 nm; GCaMP8f, 470 nm) and 2 six-port fluorescence DORIC minicubes, each with the following configuration: in nm; isosbestic excitation filters, 400–410; GCaMP excitation filter, 460–490; GCaMP emission filter, 500–540; modulated at 10 μW (isosbestic) and 30 μW (GCaMP8f), retrieved at a 1 kHz sampling rate. Behavior recordings were synchronized with fiber photometry recordings collected via DORIC Neuroscience Studio.

#### Optogenetic stimulation of OXT fibers during home cage social investigation

To study the effects of optogenetic stimulation of OXT fibers in the NAcSh or CeL on the social behavior of KO mice, we used a 3-day repeated protocol. All mice in the optogenetics cohorts were dual implanted in the NAcSh and CeL to allow simultaneous recordings from both regions. On Day1, we conducted a home cage social investigation exactly as described above, while recording from the NAcSh and CeL. On Day 2, the assay was repeated with novel stimulus mice. During the first 5 min of the investigation phase, a 555 nm laser (MGL-FN-561-100mW, UltraLasers) was delivered at 30 Hz with 10 mW intensity to either the NAcSh or the CeL to drive calcium influx in OXT fiber terminals and elicit activity-dependent OXT release. On Day 3, the same protocol was repeated using a novel stimulus mouse, with stimulation applied to the region not targeted on Day 2. GCaMP8f recordings were collected for the full 10 min from the non-stimulated region and for the final 5 min of the assay from the stimulated region.

#### Posthoc Histology

To confirm virus expression and optic fiber targeting, we transcardially perfused mice with phosphate buffer saline and 4% paraformaldehyde using previously described protocols ^115^. Brains were extracted, then sliced on a Leica VT1200 Semi-Automatic Vibrating Blade Microtome. We collected 50µm thick representative sections from all targeted regions (NAcSh, CeL, PVN). Sections were immunostained collectively to label OXT neurons and fibers and enhance GCaMP8f and ChrimsonR fluorescent reporter signal (Primary antibodies (1:2000): xNeurophysin-Rb, xGFP-Chk, xRFP-Gt; Donkey-host secondary antibodies (1:500): AlFl+ xRb-647, AlFl+ xChk-488, AlFl+ xGt-555). Immunolabeled slices were mounted on glass slides using Fluoromount with DAPI stain for nuclear labeling and coverslipped. Mounted sections were imaged 10× and 0.75 pixel/μm using Zeiss Axioscan 7 slide scanner at CALM. To confirm colocalization of OXT immunolabeled signal and ChrimsonR virus labeling of OXT fibers, ROIs were imaged using a Nikon A1R inverted confocal microscope with a 60x objective at CALM and analyzed using FIJI^116^.

### Analyses

#### Atlas registration and c-Fos+ cell counting

For tissue section images taken with a slidescanner, NeuroInfo (MBF Biosciences) was used to parcellate whole-section images of representative brain slices to the Allen Mouse Brain atlas, followed by automated detection of c-Fos+ cell density in each ROI. For whole brain images, Neuroinfo (MBF) was used for automated detection of c-Fos+ cells in thresholded images (**Fig.1**). This was followed by 2D or whole brain 3D volume parcellation in Neuroinfo using machine-learning-assisted linear registration (2D and 3D) and Symmetric Normalization (non-linear; 3D) alignment of the image to the Allen Mouse Brain Atlas. Using these analysis pipelines, c-Fos-positive cell counts were extracted for every region parcellated under the Allen Common Coordinate Framework (Allen ccf.)^117^. Cell count data were extracted from SSN regions, defined as ILA (infralimbic area), PL (prelimbic area), ACA (anterior cingulate area), PVN (hypothalamic paraventricular nucleus), NAc (nucleus accumbens), VTA (ventral tegmental area), MeA (medial amygdala), BLA (basolateral amygdala), and AON (anterior olfactory nucleus)^36,118^. We also included in our analysis additional regions that have established roles in social salience processing including HPF (hippocampal formation)^119–121^, CeA (central amygdala)^122,123^, SS (somatosensory cortex)^124^, VIS (visual cortex)^125,126^, and AUD (auditory cortex)^127,128^. Furthermore, the HPF^26,129,130^, CeA^107,131^, SS^132^, VIS^125,126^, and AUD^133–135^ are strongly associated with social behavior phenotypes in autism. Subsequent data plotting and statistical analyses were conducted using R Studio and Graphpad PRISM (v10).

#### Behavior and Fiber Photometry analysis

Behavior was manually annotated using BORIS^100^. To plot social event-triggered traces, behavior event start times (>1s long) were isolated from BORIS scoring data using custom R-scripts. Event-triggered GCAMP signal traces 10s before and 10s after the event start were extracted using an open-source MATLAB photometry analysis pipeline (FPAv.2) by Leo Molina^136^, as described in our previous study^61^. Z-scores obtained from FPAv.2 were fitted to a polynomial curve for baseline correction using custom R-scripts. Mean and standard error of the mean were plotted and processed for AUC and peak analysis using GraphPad Prism (version 10.3.1). Event-triggered GCaMP8f signal traces 10s before and 10s after the event start were extracted using an open-source MATLAB photometry analysis pipeline (FPAv.2) by Leo Molina^136^, as described in our previous study^61^. Z-scores obtained from FPAv.2 were fitted to a polynomial curve for baseline correction using custom R-scripts. Mean and standard error of the mean were plotted and processed for AUC and peak analysis using GraphPad Prism (version 10.3.1).

#### Statistical Analyses

Statistical analyses were conducted using PRISM Graphpad (v.10.5.0) and with significance thresholds set at p < .05 or q < .05 for all tests. We compared total duration of social investigation, total number of social bouts, and mean duration of individual social bouts between KO mice and their WT littermates in a sex-stratified manner using a two-way ANOVA followed by an FDR correction for multiple comparisons. To compare behavior scores across treatment (control/DREADD) conditions, we used two sample unpaired t-tests (**Fig.2**). For SSN-wide comparisons of c-Fos counts between treatment conditions across brain regions, we used a Two-way ANOVA with an FDR correction for multiple comparisons (**Fig.3**). For comparing network activity between non-social baseline and social groups or between treatment conditions, we conducted a Pearson cross-correlation analysis with data represented as a covariance matrix. We also analyzed c-Fos-count/mean social duration correlation by plotting regional ROI c-Fos counts from whole SSN datasets with mean duration of social investigation and fitting the datapoints to a linear regression line. For standardized analysis of c-Fos+ cell densities in PVN, NAc, CeA, and ILA quantified from tissue sections, we normalized all c-Fos-counts to the mean of the non-social WT group of own sex, then compared c-Fos+ cell density in each brain region across groups using a two-way ANOVA with an FDR post-hoc correction. To examine group differences in GCaMP8f and/or GRAB-OT sensor signal, we calculated maximum peak values and area under the curve (AUC) for 10 seconds before and 10 seconds after the beginning of social events. We confirmed that AUC values for treatment effect groups in the optogenetics cohorts were non-zero using a one-sample t-test. Peak values and AUCs were compared across groups using an unpaired two sample t-test. Complete summaries for statistical tests used in each figure are available in the corresponding table in the supplementary material.

## Acknowledgments

We thank C. Bourque and members of the Choe lab for their valuable feedback on the manuscript. We thank J. Renström for help with preparing tissue for post-mortem processing. We also acknowledge M. Matthews, C. McRae, I. Wang, K. Hossain, Y. K. Lee, and undergraduate research assistants in the Choe lab for mouse colony management support and genotyping. Additionally, we thank D. Graham and other McMaster CAF animal care staff for animal care assistance. Confocal and slidescanner microscopy made use of instrumentation available at the Centre for Advanced Light Microscopy (CALM) at McMaster. This study was supported by the Canada Foundation for Innovation (40750), Ontario Research Fund, Canada Research Chair (CRC-2020-00071), NSERC Discovery Grant (RGPIN-2021-03732), The Azrieli Foundation Science Grant, Canadian Institutes of Health Research Project Grant (PJT-183808), and McMaster Startup fund to KYC and the Synergy ERC grant “OxytocINspace” 101071777, German-Israeli Project (DIP) GR3619-1 and ERANET-Neuron GR 3619/25-1 to VG.

