## Supplemental materials for "Autism-risk gene mutations convergently disrupt sexually dimorphic oxytocin circuits to lower social engagement"

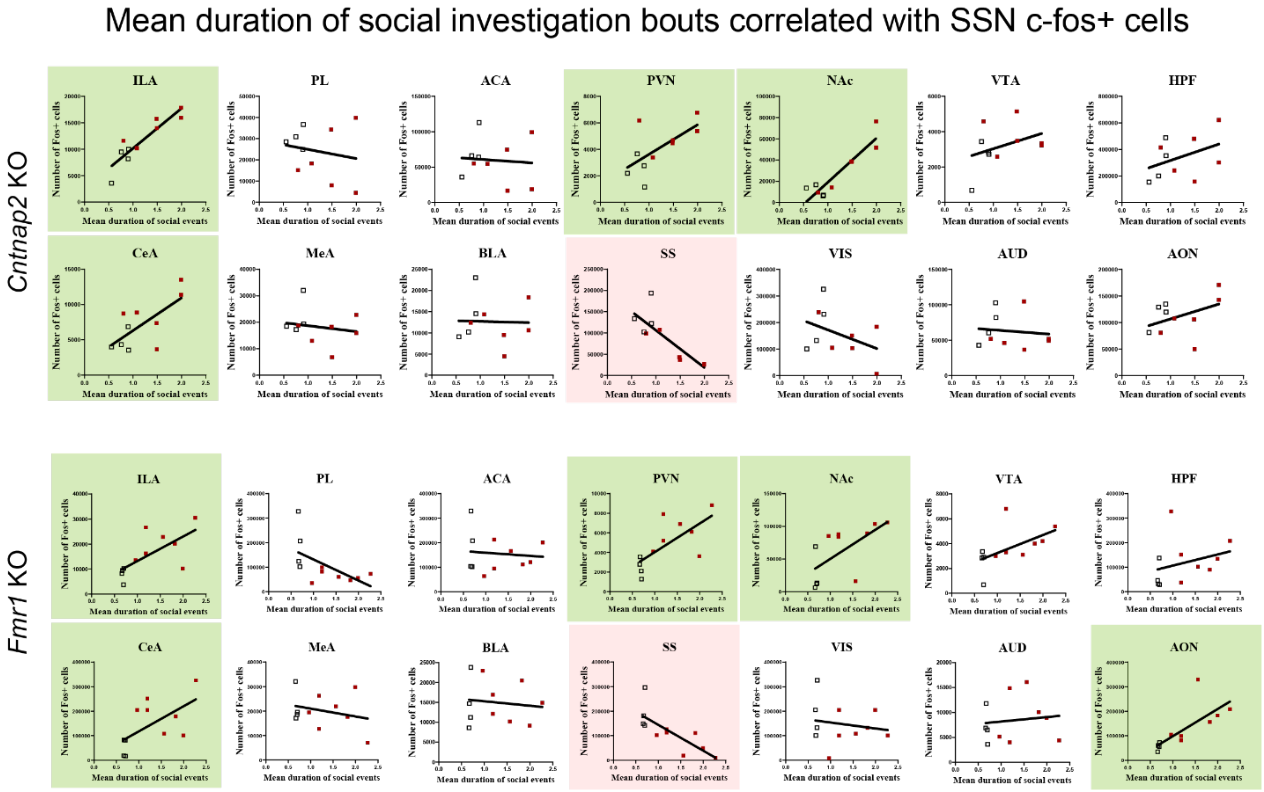


**Supplementary Figure 1.** Mean duration of spontaneous social interaction bouts correlated with cfos+ cell counts in individual SSN regions showing statistically significant (*P<0.05) strong positive correlations in ILA, PVN, NAc, and CeA (green), and negative correlation in SS (pink), consistent across genotypes (*Cntnap2* KO: top, *Fmr1* KO: bottom). Each data point represents one animal, injected with either hM3Dq or Venus (control). Square data points represent males, triangle data points represent females.

Number of social investigation bouts correlated with SSN c-fos+ cells

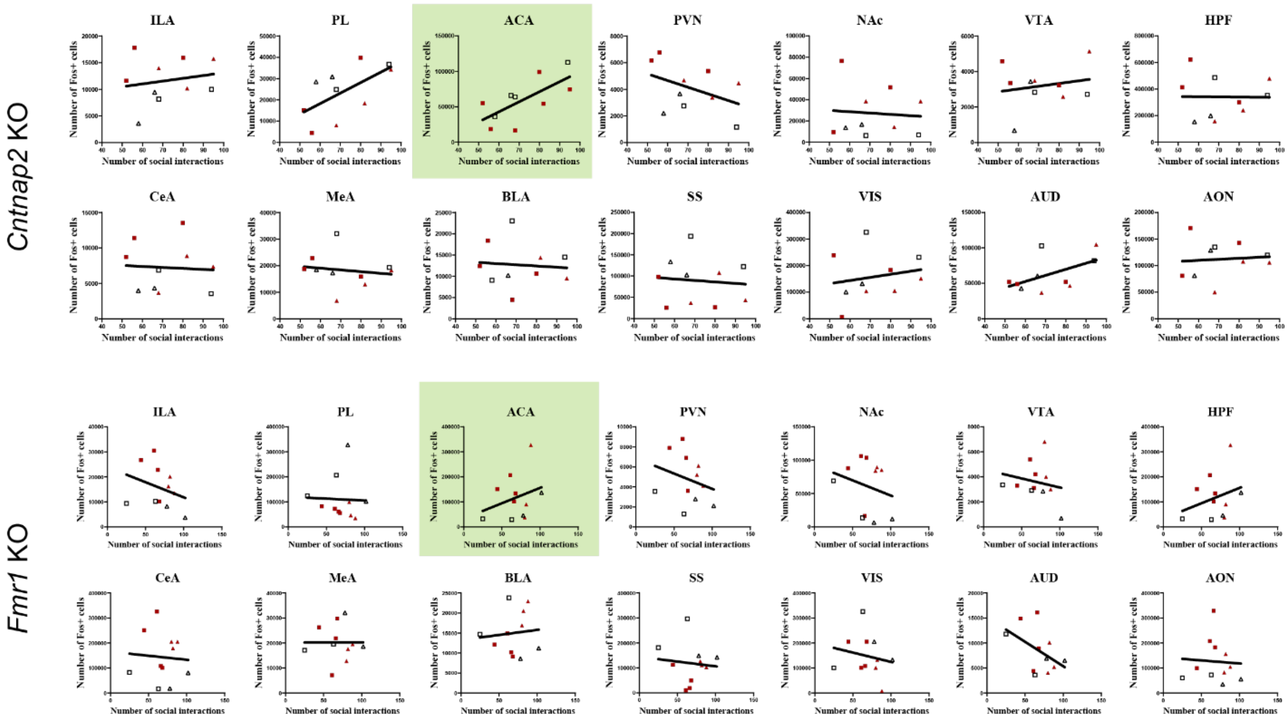


**Supplementary Figure 2.** Total number of spontaneous social investigation bouts (in 10-min social investigation period) correlated with cfos+ cell counts in individual SSN regions showing statistically significant (*P<0.05) strong positive correlation in ACA only (green), consistent across genotypes (*Cntnap2* KO: top, *Fmr1* KO: bottom). Each data point represents one animal, injected with either hM3Dq or Venus (control). Square data points represent males, triangle data points represent females.

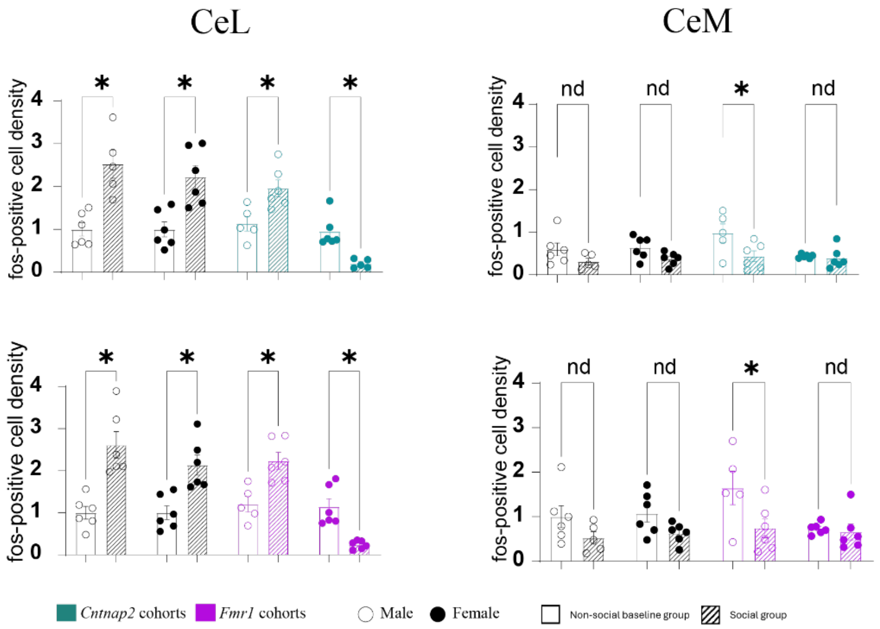


**Supplementary Fig 3.** WT baseline-normalized c-Fos+ cell density changes in the CeL in *Cntnap2* (green) and *Fmr1* KO (purple) mice and their WT littermates (black). *P<0.05.

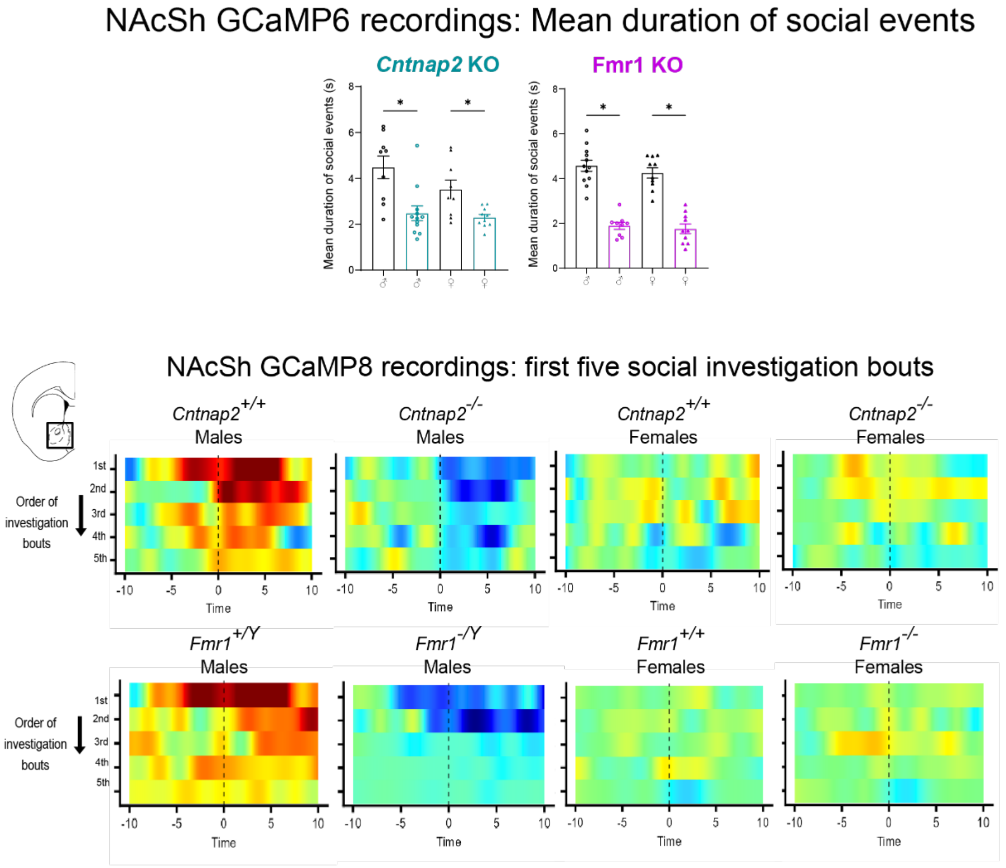


**Supplementary Fig 4.** Bar graphs**:** Sex stratified comparisons of mean duration of social investigation bouts between *Cntnap2* (green) and *Fmr1* KO (purple) mice and their WT littermates (black). Two-way ANOVA with FDR correction for multiple comparisons. *P<0.05. NAcSh GCAMP8 recording heatmaps showing averaged GCAMP8 signal (Δf/f) during the first 5 investigation bouts in male and female *Cntnap2* and *Fmr1* KO mice and their WT littermates.

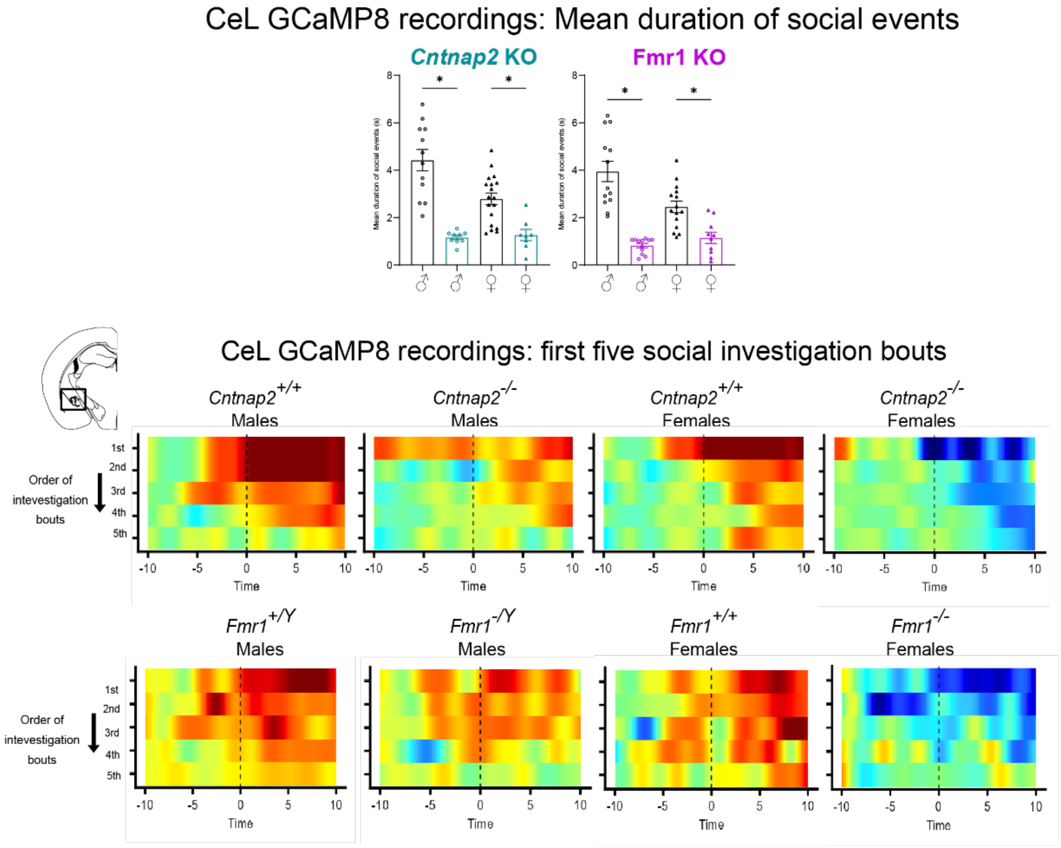


**Supplementary Fig 5.** Bar graphs**:** Sex stratified comparisons of mean duration of social investigation bouts between *Cntnap2* (green) and *Fmr1* KO (purple) mice and their WT littermates (black). Two-way ANOVA with FDR correction for multiple comparisons. *P<0.05. CeL GCAMP8 recording heatmaps showing averaged GCAMP8 signal (Δf/f) during the first 5 investigation bouts in male and female *Cntnap2* and *Fmr1* KO mice and their WT littermates.

Table1

| Analysis | | F (DFn, DFd) | P value | P value summary |
|---|---|---|---|---|
| 2-way ANOVA. *Cntnap2* cohort comparisons of Total investigation time | Interaction | F (1, 24) = 1.544 | P=0.2260 | ns |
| | Sex | F (1, 24) = 14.60 | P=0.0008 | *** |
| | Genotype | F (1, 24) = 16.00 | P=0.0005 | *** |
| 2-way ANOVA. *Fmr1* cohort comparisons of Total investigation time | Interaction | F (1, 24) = 10.60 | P=0.0034 | ** |
| | Sex | F (1, 24) = 0.6103 | P=0.4423 | ns |
| | Genotype | F (1, 24) = 23.11 | P<0.0001 | **** |
| 2-way ANOVA. *Cntnap2* cohort comparisons of Total number of social bouts | Interaction | F (1, 22) = 0.002414 | P=0.9613 | ns |
| | Sex | F (1, 22) = 0.2360 | P=0.6319 | ns |
| | Genotype | F (1, 22) = 1.310 | P=0.2647 | ns |
| 2-way ANOVA. *Fmr1* cohort comparisons of Total number of social bouts | Interaction | F (1, 22) = 6.865 | P=0.0156 | * |
| | Sex | F (1, 22) = 4.915 | P=0.0373 | * |
| | Genotype | F (1, 22) = 0.9264 | P=0.3463 | ns |
| 2-way ANOVA. *Cntnap2* cohort comparisons of Mean duration of individual social bouts | Interaction | F (1, 22) = 1.489 | P=0.2353 | ns |
| | Sex | F (1, 22) = 13.71 | P=0.0012 | ** |
| | Genotype | F (1, 22) = 51.25 | P<0.0001 | **** |
| 2-way ANOVA. *Fmr1* cohort comparisons of Mean duration of individual social bouts | Interaction | F (1, 22) = 14.32 | P=0.0010 | ** |
| | Sex | F (1, 22) = 0.8196 | P=0.3751 | ns |
| | Genotype | F (1, 22) = 46.18 | P<0.0001 | **** |

| Analysis | | t, df | P value | P value summary |
|---|---|---|---|---|
| Welch's two-sample t-test. *Cntnap2* cohort comparisons of social scores between control and treatment groups | Total investigation time | t=2.418, df=11.54 | 0.0332 | * |
| | Mean duration of individual social bouts | t=3.622, df=6.726 | 0.0091 | ** |
| | Total number of social bouts | t=1.191, df=12.08 | 0.2567 | ns |
| | Inter-event interval | t=2.328, df=12.91 | 0.0368 | * |
| Welch's two-sample t-test. *Fmr1* cohort comparisons of social scores between control and treatment groups | Total duration | t=2.416, df=9.914 | 0.0365 | * |
| | Mean duration of individual social bouts | t=3.491, df=7.357 | 0.0093 | ** |
| | Total number of social bouts | t=0.8541, df=9.121 | 0.4149 | ns |
| | Inter-event interval | t=3.117, df=9.540 | 0.0115 | * |

Table2a

| Analysis | F (DFn, DFd) | P value | P value summary |
|---|---|---|---|
| One-way ANOVA. *Cntnap2* cohort comparisons of Fos+ cell density across treament conditions and SSN regions | 3.594 (27, 112) | P<0.0001 | * |
| False Discovery Rate correction | | | |
| Dreadd - Control | Individual P Value | Individual q Value | Discovery? |
| ILA | 0.8358 | 0.8695 | No |
| PL | 0.7403 | 0.8695 | No |
| ACA | 0.9304 | 0.8695 | No |
| PVN | 0.3786 | 0.8695 | No |
| NAc | 0.9662 | 0.8695 | No |
| VTA | <0.0001 | 0.8695 | Yes |
| HPF | 0.8907 | <0.0001 | No |
| CeA | 0.7453 | 0.8695 | No |
| MeA | 0.8523 | 0.8695 | No |
| BLA | 0.0017 | 0.8695 | Yes |
| SS | 0.0346 | 0.0105 | No |
| VIS | 0.6216 | 0.1454 | No |
| AUD | 0.8323 | 0.8695 | No |
| AON | 0.7085 | 0.8695 | No |

| Analysis | F (DFn, DFd) | P value | P value summary |
|---|---|---|---|
| One-way ANOVA. *Fmr1* cohort comparisons of Fos+ cell density across treament conditions and SSN regions | 2.776 (27, 112) | P<0.0001 | * |
| False Discovery Rate correction | | | |
| Dreadd - Control | Individual P Value | Individual q Value | Discovery? |
| ILA | 0.9374 | 0.7952 | No |
| PL | 0.9199 | 0.7952 | No |
| ACA | 0.263 | 0.7595 | No |
| PVN | 0.9577 | 0.7952 | No |
| NAc | 0.621 | 0.7952 | No |
| VTA | 0.9282 | 0.7952 | No |
| HPF | <0.0001 | <0.0001 | Yes |
| CeA | 0.9291 | 0.7952 | No |
| MeA | 0.9182 | 0.7952 | No |
| BLA | 0.9639 | 0.7952 | No |
| SS | 0.0051 | 0.0197 | Yes |
| VIS | 0.0005 | 0.003 | Yes |
| AUD | 0.7893 | 0.7952 | No |
| AON | 0.9088 | 0.7952 | No |

### Table2b

**p-values for cross correlation matrix exmaining the coordinated activity of SSN regions during social behavior in *Cntnap2* KO mice**

| Control (Venus) | hM3Dq: ACA | PL | ILA | PVN | NAc | VTA | HPF | CeA | MeA | BLA | VIS | AUD | AON | SS |
|---|---|---|---|---|---|---|---|---|---|---|---|---|---|---|
| ACA | | 6.76E-05 | 0.676529 | 0.288664 | 0.344173 | 0.344733 | 0.288459 | 0.628231 | 0.968808771 | 0.404614258 | 0.051333591 | 0.145714159 | 0.751586 | 0.548341 |
| PL | 0.02479 | | 0.980836 | 0.160443 | 0.58676 | 0.291258 | 0.480634 | 0.890342 | 0.807627056 | 0.23633481 | 0.156107959 | 0.050053846 | 0.857023 | 0.331627 |
| ILA | 0.012589 | 0.232692 | | 0.254421 | 0.000162 | 0.961789 | 0.826364 | 0.605023 | 0.196456723 | 0.498251629 | 0.074798292 | 0.5045674 | 0.157854 | 0.220494 |
| PVN | 0.174625 | 0.148582 | 0.923459 | | 0.236738 | 0.954159 | 0.245433 | 0.684082 | 0.046858546 | 0.118512091 | 0.656565034 | 0.376278431 | 0.514 | 0.944736 |
| NAc | 0.179137 | 0.957198 | 0.510606 | 0.098416 | | 0.426155 | 0.822834 | 0.97716 | 0.2732159 | 0.259688302 | 0.008097901 | 0.952401077 | 0.183346 | 0.459167 |
| VTA | 0.137887 | 0.647697 | 0.000572 | 0.362866 | 0.795344 | | 0.169658 | 0.056867 | 0.508412151 | 0.348288228 | 0.136072209 | 0.016369093 | 0.722036 | 0.417808 |
| HPF | 0.267084 | 0.620127 | 0.233219 | 0.635414 | 0.003512 | 0.263079 | | 0.896659 | 0.083780459 | 0.552696614 | 0.257588025 | 0.426536171 | 0.869856 | 0.872821 |
| CeA | 0.54012 | 0.014465 | 0.902136 | 0.32425 | 0.299141 | 0.524533 | 0.045456 | | 0.661858315 | 0.085364638 | 0.603961219 | 0.463247217 | 0.619733 | 0.201968 |
| MeA | 0.898129 | 0.074737 | 0.851887 | 0.861102 | 0.056145 | 0.655075 | 0.005084 | 0.000454 | | 0.013750445 | 0.601980821 | 0.498891111 | 0.025136 | 0.87231 |
| BLA | 0.551157 | 0.311894 | 0.407817 | 0.880713 | 0.012856 | 0.354376 | 2.86E-05 | 0.008418 | 0.000252238 | | 0.167548173 | 0.54145365 | 0.069735 | 0.276487 |
| VIS | 0.281104 | 0.597052 | 0.238117 | 0.658125 | 0.004021 | 0.262026 | 3.67E-12 | 0.041028 | 0.004409621 | 1.9226E-05 | | 0.626654726 | 0.181951 | 0.866485 |
| AUD | 0.19438 | 0.724694 | 0.116231 | 0.773424 | 0.011589 | 0.128121 | 4.69E-06 | 0.056795 | 0.012790722 | 0.000248809 | 4.75998E-06 | | 0.273848 | 0.387666 |
| AON | 0.193904 | 0.961789 | 0.005115 | 0.420293 | 0.41738 | 0.000295 | 0.060678 | 0.179886 | 0.224582104 | 0.08466547 | 0.059402483 | 0.0200333 | | 0.825726 |
| SS | 0.461017 | 0.034273 | 0.612227 | 0.024838 | 0.975978 | 0.195643 | 0.259864 | 0.004544 | 0.065128889 | 0.12847476 | 0.245059728 | 0.212178142 | 0.078685 | |

**p-values for cross correlation matrix exmaining the coordinated activity of SSN regions during social behavior in *Fmr1* KO mice**

| Control (Venus) | hM3Dq: ACA | PL | ILA | PVN | NAc | VTA | HPF | CeA | MeA | BLA | VIS | AUD | AON | SS |
|---|---|---|---|---|---|---|---|---|---|---|---|---|---|---|
| ACA | | 1.39E-05 | 0.409442 | 0.487847 | 0.117213 | 0.341824 | 0.480657 | 0.949563 | 0.857085964 | 0.453814356 | 0.04260597 | 0.43578931 | 0.48726 | 0.182932 |
| PL | 0.024534 | | 0.393976 | 0.459572 | 0.084972 | 0.179709 | 0.37554 | 0.722305 | 0.524570568 | 0.318575954 | 0.051024578 | 0.270270398 | 0.320355 | 0.153318 |
| ILA | 0.005952 | 0.268017 | | 1.28E-10 | 0.966147 | 0.619852 | 0.113093 | 0.012201 | 0.582687877 | 0.009243347 | 0.17173719 | 0.506502111 | 0.001547 | 0.01917 |
| PVN | 0.16791 | 0.087096 | 0.864737 | | 0.931934 | 0.682024 | 0.081553 | 0.009514 | 0.60305481 | 0.009431026 | 0.184123418 | 0.426419823 | 0.00103 | 0.029658 |
| NAc | 0.174674 | 0.831992 | 0.400526 | 0.195306 | | 0.131339 | 0.830148 | 0.578736 | 0.030974504 | 0.28625213 | 0.007206028 | 0.818301901 | 0.324819 | 0.293872 |
| VTA | 0.078949 | 0.710091 | 8.78E-05 | 0.339679 | 0.552032 | | 0.359567 | 0.114761 | 0.029983624 | 0.87902962 | 0.610999543 | 0.032161296 | 0.399658 | 0.020514 |
| HPF | 0.615585 | 0.084282 | 0.414383 | 0.549307 | 0.019516 | 0.476016 | | 0.179866 | 0.875255404 | 0.644296589 | 0.444307071 | 0.000793625 | 0.086192 | 0.976236 |
| CeA | 0.507564 | 0.004186 | 0.877458 | 0.219313 | 0.15912 | 0.475706 | 0.031798 | | 0.268786475 | 0.137076284 | 0.352825608 | 0.859903384 | 0.000662 | 0.03157 |
| MeA | 0.852229 | 0.032917 | 0.826631 | 0.685002 | 0.021375 | 0.574112 | 0.003434 | 8.4E-05 | | 0.349950085 | 0.55557016 | 0.497783356 | 0.526619 | 0.695754 |
| BLA | 0.580885 | 0.180707 | 0.373555 | 0.917612 | 0.003132 | 0.269107 | 0.016854 | 0.002518 | 4.74316E-05 | | 0.875892943 | 0.863420176 | 0.09467 | 0.24564 |
| VIS | 0.50736 | 0.214533 | 0.295972 | 0.892978 | 0.003452 | 0.206733 | 0.027471 | 0.003368 | 0.000139072 | 7.44949E-10 | | 0.438591421 | 0.035908 | 0.143157 |
| AUD | 0.506234 | 0.221596 | 0.324433 | 0.96223 | 0.002289 | 0.239795 | 0.021125 | 0.004036 | 0.000122835 | 8.52341E-11 | 4.6181E-11 | | 0.402182 | 0.366503 |
| AON | 0.354765 | 0.149135 | 0.092446 | 0.331671 | 0.112201 | 0.092069 | 0.000212 | 0.200537 | 0.072865857 | 0.18499162 | 0.242305723 | 0.207981795 | | 0.045483 |
| SS | 0.651712 | 0.185463 | 0.586493 | 0.815712 | 0.000853 | 0.502613 | 0.003153 | 0.005571 | 4.46537E-05 | 1.75801E-06 | 9.89457E-06 | 2.97788E-06 | 0.072209 | |

**Mean duration of social investigation bouts correlated with the number of fos+cells *Cntnap2* KO mice**

| Region | ACA | PL | ILA | PVN | NAc | VTA | HPF | CeA | MeA | BLA | VIS | AUD | AON | SS |
|---|---|---|---|---|---|---|---|---|---|---|---|---|---|---|
| F | 0.04933 | 0.2803 | 37.57 | 5.837 | 42.48 | 1.284 | 1.552 | 7.311 | 0.2594 | 0.006869 | 1.51 | 0.09573 | 1.65 | 16.51 |
| DFn, DFd | 1, 8 | 1, 8 | 1, 8 | 1, 8 | 1, 8 | 1, 8 | 1, 8 | 1, 8 | 1, 8 | 1, 8 | 1, 8 | 1, 8 | 1, 8 | 1, 8 |
| P value | 0.8298 | 0.6109 | 0.0003 | 0.0421 | 0.0002 | 0.2899 | 0.2481 | 0.0269 | 0.6243 | 0.936 | 0.2541 | 0.7649 | 0.235 | 0.0036 |
| P value summary | NS | NS | S | S | S | NS | NS | S | NS | NS | NS | NS | NS | S |

**Mean duration of social investigation bouts correlated with the number of fos+cells in *Fmr1* KO mice**

| Region | ACA | PL | ILA | PVN | NAc | VTA | HPF | CeA | MeA | BLA | VIS | AUD | AON | SS |
|---|---|---|---|---|---|---|---|---|---|---|---|---|---|---|
| F | 0.2361 | 1.261 | 7.581 | 8.586 | 6.298 | 3.728 | 0.8542 | 5.333 | 0.6472 | 0.1325 | 0.266 | 0.1285 | 11.12 | 12.85 |
| DFn, DFd | 1, 9 | 1, 9 | 1, 9 | 1, 9 | 1, 9 | 1, 9 | 1, 9 | 1, 9 | 1, 9 | 1, 9 | 1, 9 | 1, 9 | 1, 9 | 1, 9 |
| P value | 0.6387 | 0.2904 | 0.0224 | 0.0168 | 0.0333 | 0.0856 | 0.3795 | 0.0463 | 0.4419 | 0.7243 | 0.6185 | 0.7282 | 0.0087 | 0.0059 |
| P value summary | NS | NS | S | S | S | NS | NS | S | NS | NS | NS | NS | S | S |

**Total number of of social investigation bouts correlated with the number of fos+cells *Cntnap2* KO mice**

| Region | ACA | PL | ILA | PVN | NAc | VTA | HPF | CeA | MeA | BLA | VIS | AUD | AON | SS |
|---|---|---|---|---|---|---|---|---|---|---|---|---|---|---|
| F | 7.334 | 4.927 | 0.2897 | 1.839 | 0.0529 | 0.3239 | 0.0007889 | 0.03086 | 0.1914 | 0.05706 | 0.3238 | 3.135 | 0.05638 | 0.07519 |
| DFn, DFd | 1, 8 | 1, 8 | 1, 8 | 1, 8 | 1, 8 | 1, 8 | 1, 8 | 1, 8 | 1, 8 | 1, 8 | 1, 8 | 1, 8 | 1, 8 | 1, 8 |
| P value | 0.0267 | 0.0572 | 0.605 | 0.2121 | 0.8239 | 0.5849 | 0.9783 | 0.8649 | 0.6733 | 0.8172 | 0.585 | 0.1146 | 0.8183 | 0.7909 |
| P value summary | S | NS | NS | NS | NS | NS | NS | NS | NS | NS | NS | NS | NS | NS |

**Total number of of social investigation bouts correlated with the number of fos+cells *Fmr1* KO mice**

| Region | ACA | PL | ILA | PVN | NAc | VTA | HPF | CeA | MeA | BLA | VIS | AUD | AON | SS |
|---|---|---|---|---|---|---|---|---|---|---|---|---|---|---|
| F | 47.8 | 0.02161 | 0.9097 | 0.6901 | 0.5499 | 0.3639 | 0.1918 | 0.04376 | 0.00003261 | 0.08837 | 0.3352 | 2.602 | 0.03249 | 0.09357 |
| DFn, DFd | 1, 9 | 1, 9 | 1, 9 | 1, 9 | 1, 9 | 1, 9 | 1, 9 | 1, 9 | 1, 9 | 1, 9 | 1, 9 | 1, 9 | 1, 9 | 1, 9 |
| P value | <0.0001 | 0.8864 | 0.3651 | 0.4276 | 0.4773 | 0.5613 | 0.6718 | 0.839 | 0.9956 | 0.773 | 0.5768 | 0.1412 | 0.8609 | 0.7666 |
| P value summary | S | NS | NS | NS | NS | NS | NS | NS | NS | NS | NS | NS | NS | NS |

### Table3

**p-values for cross correlation matrix exmaining the coordinated activity of the ILA, PVN, Nac and CeA during social behavior in *Cntnap2* $^{+/+}$ mice**

| | | home cage + social | | | |
|---|---|---|---|---|---|
| | | ILA | PVN | NAc | CeA |
| home cage | ILA | | 0.000352378 | 0.003690323 | 0.007182105 |
| | PVN | 0.370172498 | | 0.000294215 | 0.000287345 |
| | NAc | 0.716432132 | 0.911866938 | | 0.000317953 |
| | CeA | 0.665675622 | 0.44558642 | 0.586673311 | |

**p-values for cross correlation matrix exmaining the coordinated activity of the ILA, PVN, Nac and CeA during social behavior in *Cntnap2* $^{-/-}$ mice**

| | | home cage + social | | | |
|---|---|---|---|---|---|
| | | ILA | PVN | NAc | CeA |
| home cage | ILA | | 0.015504082 | 0.928142284 | 0.449479799 |
| | PVN | 0.619300365 | | 0.819561832 | 0.105250842 |
| | NAc | 0.755748699 | 0.248445243 | | 0.752791142 |
| | CeA | 0.696275874 | 0.380902933 | 0.087188084 | |

**p-values for cross correlation matrix exmaining the coordinated activity of the ILA, PVN, Nac and CeA during social behavior in *Fmr1* $^{+/+}$ mice**

| | | home cage + social | | | |
|---|---|---|---|---|---|
| | | ILA | PVN | NAc | CeA |
| home cage | ILA | | 0.045265089 | 0.0182996 | 0.030731047 |
| | PVN | 0.572672596 | | 0.000129937 | 0.0004692 |
| | NAc | 0.832385305 | 0.775610277 | | 7.18183E-05 |
| | CeA | 0.406016569 | 0.104545471 | 0.463320825 | |

**p-values for cross correlation matrix exmaining the coordinated activity of the ILA, PVN, Nac and CeA during social behavior in *Fmr1* $^{-/-,-/Y}$ mice**

| | | home cage + social | | | |
|---|---|---|---|---|---|
| | | ILA | PVN | NAc | CeA |
| home cage | ILA | | 0.045265089 | 0.0182996 | 0.030731047 |
| | PVN | 0.572672596 | | 0.000129937 | 0.0004692 |
| | NAc | 0.832385305 | 0.775610277 | | 7.18E-05 |
| | CeA | 0.406016569 | 0.104545471 | 0.463320825 | |

| Analysis | | F (DFn, DFd) | P value | P value summary |
|---|---|---|---|---|
| 2-way ANOVA. *Cntnap2* cohort comparisons of fos+ cell density in the NAc at baseline versus after social interaction | Interaction | F (3, 38) = 8.716 | P=0.0002 | *** |
| | Behavioral group | F (1, 38) = 0.9389 | P=0.3387 | ns |
| | Sex/Genotype group | F (3, 38) = 2.719 | P=0.0580 | ns |
| 2-way ANOVA. *Fmr1* cohort comparisons of fos+ cell density in the NAc at baseline versus after social interaction | Interaction | F (3, 41) = 11.95 | P<0.0001 | **** |
| | Behavioral group | F (1, 41) = 1.227 | P=0.2744 | ns |
| | Sex/Genotype group | F (3, 41) = 5.103 | P=0.0043 | ** |
| 2-way ANOVA. *Cntnap2* cohort comparisons of fos+ cell density in the PVN at baseline versus after social interaction | Interaction | F (3, 81) = 6.679 | P=0.0004 | *** |
| | Behavioral group | F (1, 81) = 0.2191 | P=0.6410 | ns |
| | Sex/Genotype group | F (3, 81) = 2.923 | P=0.0389 | * |
| 2-way ANOVA. *Fmr1* cohort comparisons of fos+ cell density in the PVN at baseline versus after social interaction | Interaction | F (3, 77) = 26.35 | P<0.0001 | **** |
| | Behavioral group | F (1, 77) = 4.549 | P=0.0361 | * |
| | Sex/Genotype group | F (3, 77) = 1.484 | P=0.2256 | ns |
| 2-way ANOVA. *Cntnap2* cohort comparisons of fos+ cell density in the CeA at baseline versus after social interaction | Interaction | F (3, 38) = 1.500 | 0.23 | ns |
| | Behavioral group | F (1, 38) = 11.76 | 0.0015 | ** |
| | Sex/Genotype group | F (3, 38) = 2.474 | 0.0763 | ns |
| 2-way ANOVA. *Fmr1* cohort comparisons of fos+ cell density in the CeA at baseline versus after social interaction | Interaction | F (3, 38) = 1.407 | 0.2556 | ns |
| | Behavioral group | F (1, 38) = 11.02 | 0.002 | ** |
| | Sex/Genotype group | F (3, 38) = 2.354 | 0.0873 | ns |

Table4

| Analysis | Sex | | P value | P value summary |
|---|---|---|---|---|
| Welch's Two-tailed test. Comparing **Peak** and **AUC** values from social-induced **NAcSh GCAMP8** traces between *Cntnap2* KO mice and WT littermates | Male | Peak | P<0.0001 | **** |
| | | AUC | P<0.0001 | **** |
| Welch's Two-tailed test. Comparing **Peak** and **AUC** values from social-induced **NAcSh GCAMP8** traces between *Fmr1* KO mice and WT littermates | Male | Peak | P<0.0001 | **** |
| | | AUC | P<0.0001 | **** |
| Welch's Two-tailed test. Comparing **Peak** and **AUC** values from social-induced **GRAB-OT** traces from the **NAcSh** between *Cntnap2* KO mice and WT littermates | Male | Peak | P<0.0001 | **** |
| | | AUC | P<0.0001 | **** |
| Welch's Two-tailed test. Comparing **Peak** and **AUC** values from social-induced **GRAB-OT** traces from the **NAcSh** between *Cntnap2* KO mice and WT littermates | Female | Peak | P=0.7158 | ns |
| | | AUC | P=0.6157 | ns |
| Welch's Two-tailed test. Comparing **Peak** and **AUC** values from social-induced **GRAB-OT** traces from the **NAcSh** between *Fmr1* KO mice and WT littermates | Male | Peak | P<0.0001 | **** |
| | | AUC | P<0.0001 | **** |
| Welch's Two-tailed test. Comparing **Peak** and **AUC** values from social-induced **GRAB-OT** traces from the **NAcSh** between *Fmr1* KO mice and WT littermates | Female | Peak | P=0.8461 | ns |
| | | AUC | P=0.4592 | ns |

Table5

| Analysis | Sex | | P value | P value summary |
|---|---|---|---|---|
| Welch's Two-tailed test. Comparing **Peak** and **AUC** values from social-induced **CeL GCAMP8** traces between *Cntnap2* KO mice and WT littermates | Female | Peak | P=0.0327 | **** |
| | | AUC | P<0.0001 | * |
| Welch's Two-tailed test. Comparing **Peak** and **AUC** values from social-induced **CeL GCAMP8** traces between *Fmr1* KO mice and WT littermates | Female | Peak | P<0.0001 | **** |
| | | AUC | P<0.0001 | **** |
| Welch's Two-tailed test. Comparing **Peak** and **AUC** values from social-induced **GRAB-OT** traces from the **CeL** between *Cntnap2* KO mice and WT littermates | Male | Peak | P=0.7999 | ns |
| | | AUC | P=0.6863 | ns |
| Welch's Two-tailed test. Comparing **Peak** and **AUC** values from social-induced **GRAB-OT** traces from the **CeL** between *Cntnap2* KO mice and WT littermates | Female | Peak | P<0.0001 | **** |
| | | AUC | P<0.0001 | **** |
| Welch's Two-tailed test. Comparing **Peak** and **AUC** values from social-induced **GRAB-OT** traces from the **CeL** between *Fmr1* KO mice and WT littermates | Male | Peak | P=0.9739 | ns |
| | | AUC | P=7886 | ns |
| Welch's Two-tailed test. Comparing **Peak** and **AUC** values from social-induced **GRAB-OT** traces from the **CeL** between *Fmr1* KO mice and WT littermates | Female | Peak | P<0.0001 | **** |
| | | AUC | P<0.0001 | **** |

Table6a

| Analysis | | F (DFn, DFd) | P value | P value summary |
|---|---|---|---|---|
| 2-way repeated measures ANOVA. *Cntnap2* KO mouse comparisons of Mean duration of social interactions before and after optogenetic NAcSh treatment | Interaction | F (1, 4) = 12.33 | P=0.0246 | * |
| | Treatment | F (1, 4) = 17.12 | P=0.0144 | * |
| | Sex | F (1, 4) = 19.24 | P=0.0118 | * |
| | Subject | F (4, 4) = 1.081 | P=0.4709 | ns |
| Two-stage linear step-up procedure of Benjamini, Krieger and Yekutieli (FDR) | | | q value | q value summary |
| | | Male | 0.0059 | yes |
| | | Female | 0.3573 | no |
| 2-way repeated measures ANOVA. *Fmr1* KO mouse comparisons of Mean duration of social interactions before and after optogenetic NAcSh treatment | Interaction | F (1, 7) = 8.656 | P=0.0216 | * |
| | Treatment | F (1, 7) = 11.42 | P=0.0118 | * |
| | Sex | F (1, 7) = 9.954 | P=0.0160 | * |
| | Subject | F (7, 7) = 0.4598 | P=0.8365 | ns |
| Two-stage linear step-up procedure of Benjamini, Krieger and Yekutieli (FDR) | | | q value | q value summary |
| | | Male | 0.0022 | yes |
| | | Female | 0.4083 | no |
| Welch's Two-tailed test. Comparing **AUC** values from social-induced **NacSh GCAMP8** traces in *Cntnap2* KO mice between baseline and post-opto stimulation | Male | AUC | P<0.0001 | **** |
| Welch's Two-tailed test. Comparing **AUC** values from social-induced **NacSh GCAMP8** traces in *Cntnap2* KO mice between baseline and post-opto stimulation | Female | AUC | P=0.7185 | ns |
| Welch's Two-tailed test. Comparing **AUC** values from social-induced **NAcSh GCAMP8** traces in *Fmr1* KO mice between baseline and post-opto stimulation | Male | AUC | P<0.0001 | **** |
| Welch's Two-tailed test. Comparing **AUC** values from social-induced **NAcSh GCAMP8** traces in *Fmr1* KO mice between baseline and post-opto stimulation | Female | AUC | P=0.2691 | ns |

Table6b

| Analysis | | F (DFn, DFd) | P value | P value summary |
|---|---|---|---|---|
| 2-way repeated measures ANOVA. *Cntnap2* KO mouse comparisons of Mean duration of social interactions before and after optogenetic **CeL** treatment | Interaction | F (1, 4) = 21.21 | P=0.0100 | ** |
| | Treatment | F (1, 4) = 2.063 | P=0.2243 | ns |
| | Sex | F (1, 4) = 8.766 | P=0.0415 | * |
| | Subject | F (4, 4) = 1.000 | P=0.4999 | ns |
| Two-stage linear step-up procedure of Benjamini, Krieger and Yekutieli (FDR) | | | q value | q value summary |
| | | Male | 0.0465 | yes |
| | | Female | 0.0136 | yes |
| 2-way repeated measures ANOVA. *Fmr1* KO mouse comparisons of Mean duration of social interactions before and after optogenetic **CeL** treatment | Interaction | F (1, 6) = 167.2 | P<0.0001 | **** |
| | Treatment | F (1, 6) = 84.85 | P<0.0001 | **** |
| | Sex | F (1, 6) = 17.61 | P=0.0057 | ** |
| | Subject | F (6, 6) = 14.68 | P=0.0024 | ** |
| Two-stage linear step-up procedure of Benjamini, Krieger and Yekutieli (FDR) | | | q value | q value summary |
| | | Male | 0.041 | yes |
| | | Female | <0.0001 | yes |
| Welch's Two-tailed test. Comparing **AUC** values from social-induced **CeL GCAMP8** traces in *Cntnap2* KO mice between baseline and post-opto stimulation | Male | AUC | P=0.0101 | * |
| Welch's Two-tailed test. Comparing **AUC** values from social-induced **CeL GCAMP8** traces in *Cntnap2* KO mice between baseline and post-opto stimulation | Female | AUC | P=0.0023 | ** |
| Welch's Two-tailed test. Comparing **AUC** values from social-induced **CeL GCAMP8** traces in *Fmr1* KO mice between baseline and post-opto stimulation | Male | AUC | P=0.006 | ** |
| Welch's Two-tailed test. Comparing **AUC** values from social-induced **CeL GCAMP8** traces in *Fmr1* KO mice between baseline and post-opto stimulation | Female | AUC | P<0.0001 | **** |
